# Srsf3 mediates alternative RNA splicing downstream of PDGFRα signaling

**DOI:** 10.1101/2020.12.08.416461

**Authors:** Brenna J.C. Dennison, Eric D. Larson, Rui Fu, Julia Mo, Katherine A. Fantauzzo

**Affiliations:** Department of Craniofacial Biology, University of Colorado Anschutz Medical Campus, Aurora, CO 80045, USA; RNA Bioscience Initiative, University of Colorado Anschutz Medical Campus, Aurora, CO 80045, USA; Department of Otolaryngology, University of Colorado Anschutz Medical Campus, Aurora, CO 80045, USA

**Keywords:** PDGFRα, Srsf3, palatal mesenchyme, neural crest, facial clefting, alternative RNA splicing, protein kinases

## Abstract

Signaling through the platelet-derived growth factor receptor alpha (PDGFRα) is critical for mammalian craniofacial development, though the mechanisms by which the activity of downstream intracellular effectors is regulated to mediate gene expression changes have not been defined. We find that the RNA-binding protein Srsf3 is phosphorylated at Akt consensus sites downstream of PI3K-mediated PDGFRα signaling in palatal mesenchyme cells, leading to its nuclear translocation. We further demonstrate that ablation of *Srsf3* in the neural crest lineage leads to facial clefting due to defective cranial neural crest cell specification and survival. Finally, we show that Srsf3 regulates the alternative RNA splicing of transcripts encoding protein kinases in the facial process mesenchyme to negatively regulate PDGFRα signaling. Collectively, our findings reveal that PI3K/Akt-mediated PDGFRα signaling primarily modulates gene expression through alternative RNA splicing in the facial mesenchyme and identify Srsf3 as a critical regulator of craniofacial development.

## Introduction

Craniofacial development is a complex morphogenetic process that upon disruption results in several of the most prevalent birth defects in humans (Mai et al., 2019). Signaling through the receptor tyrosine kinase platelet-derived growth factor receptor alpha (PDGFRα) is critical for this process in both humans and mice. Missense mutations in the *PDGFRA* coding region and single base-pair substitutions in the 3’ untranslated region are associated with nonsyndromic cleft palate (Rattanasopha et al., 2012). Similarly, *Pdgfra* mutant mouse models display phenotypes ranging from a cleft palate to complete facial clefting (Klinghoffer et al., 2002; Soriano, 1997; Tallquist and Soriano, 2003). Phosphatidylinositol 3-kinase (PI3K) has been identified as the primary effector of PDGFRα signaling during skeletal development in the mouse (Klinghoffer et al., 2002). Activated PI3K signaling results in the recruitment of the serine/threonine kinase Akt to the cell membrane. Once phosphorylated, active Akt dissociates from the membrane and phosphorylates hundreds of target proteins with roles in diverse cellular processes (Manning and Cantley, 2007). To identify which proteins are phosphorylated by Akt downstream of PI3K-mediated PDGFRα signaling, we previously performed a phosphoproteomic screen using primary mouse embryonic palatal mesenchyme (MEPM) cells, ultimately identifying 56 proteins that were differentially phosphorylated upon PDGF-AA ligand treatment (Fantauzzo and Soriano, 2014). A gene ontology (GO) analysis of these proteins indicated that the most significant terms for biological process all related to RNA processing, including the top two terms mRNA processing (14/56 target proteins, p = 1.4 × 10^−8^) and RNA splicing (12/56 target proteins, p = 2.5 × 10^−7^) (Fantauzzo and Soriano, 2014).

Alternative RNA splicing (AS), characterized by differential inclusion of exons in a mature RNA transcript, is an important mechanism used to regulate gene expression and increase the diversity of protein isoforms (Licatalosi and Darnell, 2010; Wang et al., 2008). Approximately 95% of multi-exon human genes are subject to AS, often in a tissue-specific manner (Pan et al., 2008; Wang et al., 2008). Dysregulation of AS has been shown to cause numerous diseases, stemming from mutations in precursor RNA sequence elements that regulate splicing, mutations in core spliceosome components and/or mutations in auxiliary RNA-binding proteins (RBPs) (Scotti and Swanson, 2016). These RBPs are trans-acting factors that bind splicing regulatory elements in introns and/or exons of pre-mRNA to promote exon inclusion or skipping (Fu and Ares, 2014; Licatalosi and Darnell, 2010). AS mediated through RBPs is an essential process in the developing mouse face, as demonstrated by the midline facial clefting and craniofacial bone hypoplasia phenotypes resulting from global and/or tissue-specific ablation of *Esrp1*, *Esrp2* and *Rbfox2* (Bebee et al., 2015; Cibi et al., 2019; Lee et al., 2020). Despite these recent studies, the critical process of AS remains understudied in neural crest cells (NCCs) and their derivatives in the facial mesenchyme. Further, the question of how the activity of RBPs is regulated to affect the AS of subsets of transcripts in a tissue-specific and spatiotemporal manner has not been explored in the context of craniofacial development.

Here, we focused on one of the RBPs detected in our phosphoproteomic screen, Serine/arginine-rich splicing factor 3 (Srsf3). Srsf3 has been implicated in NCC development in non-mouse models, as overexpression of Srsf3 mRNA in *Xenopus* led to a change in the shape and location of the expression domain of the NCC marker *Slug* (Dichmann et al., 2008). Srsf3 belongs to the highly-conserved and widely-expressed family of serine/arginine-rich (SR) proteins, which generally bind exonic splicing enhancer elements to promote exon inclusion (Fu and Ares, 2014; Licatalosi and Darnell, 2010). Srsf3 specifically was shown to bind pyrimidine-rich motifs in both exons and introns in murine embryonic carcinoma cells, with a preference for the former (Änkö et al., 2012). Importantly, RBPs can be regulated by post-translational modifications, including phosphorylation, that can alter several aspects of their function, such as subcellular localization, RNA-binding and/or sequence specificity (Stamm, 2008). Phosphorylation of Akt consensus sites within the C-terminal arginine/serine-rich (RS) domain of Srsf3 has been shown to drive its translocation to the nucleus (Bavelloni et al., 2014; Long et al., 2019), though the upstream inputs that stimulate these modifications are incompletely understood.

In this work, we demonstrated that PI3K/Akt-mediated PDGFRα signaling primarily regulates the expression of genes involved in palatal shelf morphogenesis through AS, in part through the phosphorylation and subsequent nuclear translocation of Srsf3. We further showed that ablation of *Srsf3* in the NCC lineage leads to a severe midline facial clefting phenotype due to defective NCC specification and survival, and the mis-splicing of multiple protein kinases. Taken together, our results provide significant insight into the mechanisms underlying gene expression regulation during mammalian craniofacial development.

## Results

### Differential alternative splicing of select transcripts in response to PI3K-mediated PDGFRα signaling

To determine if disrupted PI3K-mediated PDGFRα signaling in the secondary palatal shelves (PS) leads to changes in AS, we harvested and sequenced PS mesenchyme RNA from three biological replicates of embryonic day 13.5 (E13.5) *Pdgfra*^*+/+*^ versus *Pdgfra*^*PI3K/PI3K*^ embryos in which PDGFRα is unable to bind PI3K (Klinghoffer et al., 2002) (Table S1). rMATS (Shen et al., 2014) was used to detect AS events, identifying 523 events that were significantly different between genotypes across the five major classes, with the majority of events (75.7%) involving skipped exons (SE) (Table S2; Figure 1A). A GO analysis of the 348 genes represented in the SE class using the MGI Mammalian Phenotype Level 4 2019 library of the Enrichr gene list enrichment analysis tool (Chen et al., 2013; Kuleshov et al., 2016) revealed a number of significant terms corresponding to phenotypes observed in *Pdgfra*^*PI3K/PI3K*^ embryos, including abnormal secondary palate development (p = 0.03).

**Figure 1.**
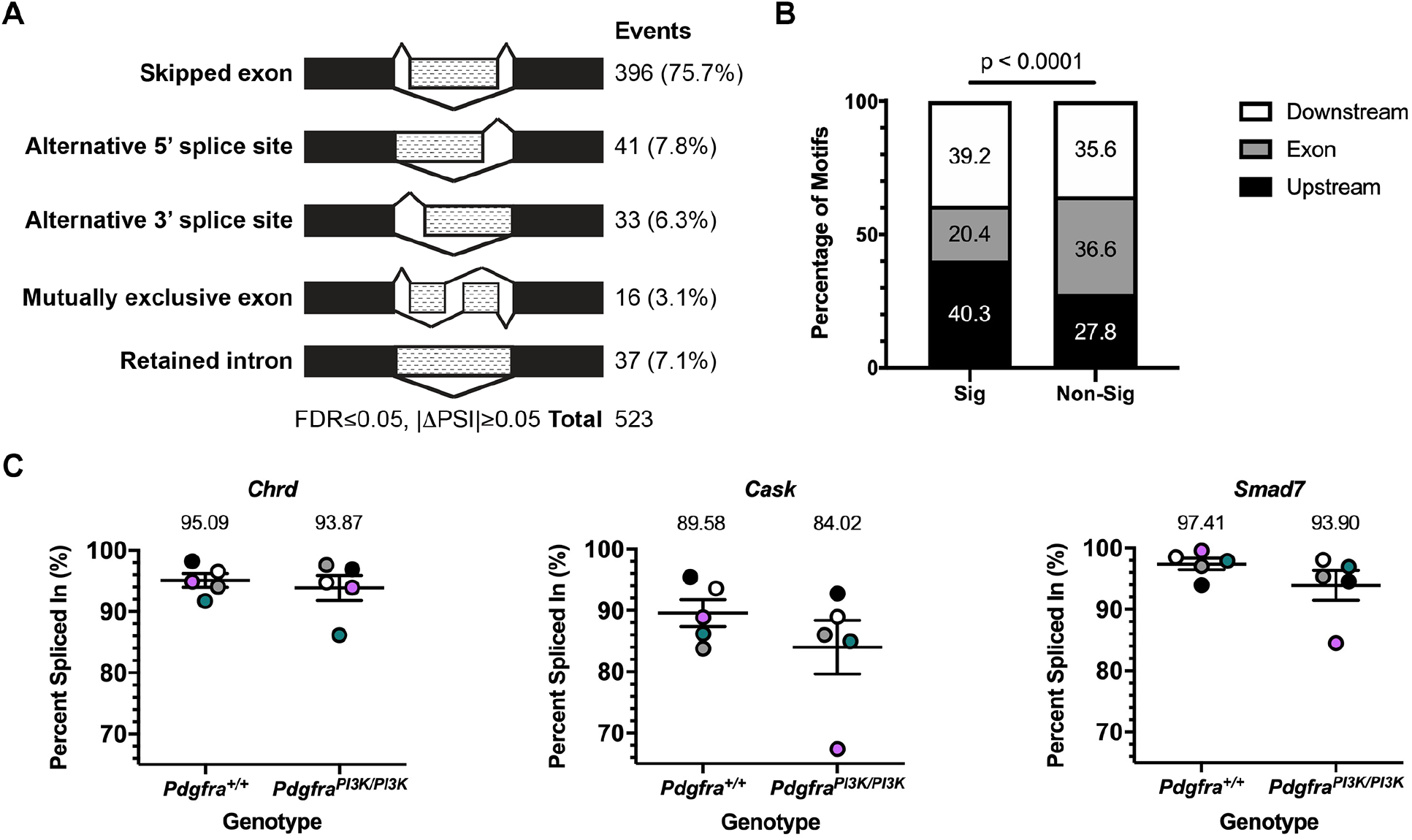
Differential alternative splicing of select transcripts in response to PI3K-mediated PDGFRα signaling. (A) Summary of differential AS events detected by rMATS analysis of RNA-seq data from E13.5 *Pdgfra*^*+/+*^ versus *Pdgfra*^*PI3K/PI3K*^ PS mesenchyme. FDR, false discovery rate; ΔPSI, difference in percent spliced in. (B) Bar graph depicting the percentage of Srsf3 motifs detected by RBPmap within or flanking exons between significant (Sig) and non-significant (Non-Sig) skipped exon events. (C) Scatter dot plots depicting percent spliced in for three differentially alternatively-spliced transcripts between E13.5 *Pdgfra*^*+/+*^ versus *Pdgfra*^*PI3K/PI3K*^ PS mesenchyme as assessed by qPCR. Data are presented as mean ± SEM. Colors correspond to independent experiments across five biological replicates of *Pdgfra*^*+/+*^ and *Pdgfra*^*PI3K/PI3K*^ samples.

We then used RBPmap (Paz et al., 2014) to examine annotated Srsf3 motifs in the 396 SE plus 250 bp flanking each end, revealing Srsf3 motifs in 381 (96.2%) of these regions of interest (ROIs) (Table S2). Though the density of these motifs was similar between significant (4.4 motifs per ROI) versus non-significant (4.6 motifs per ROI) SE events in our dataset (p = 0.2), Srsf3 motifs were enriched upstream and downstream of the SE in the significant SE events (p-value < 0.0001) (Figure 1B). Among the genes represented in the SE class that had one or more Srsf3 motifs in the ROI, 22 have a corresponding mouse model with a craniofacial phenotype, eight of which have a cleft secondary palate (Table S3). We examined the differential AS of three of these transcripts, *Chrd, Cask* and *Smad7* (Atasoy et al., 2007; Bachiller et al., 2003; Papangeli and Scambler, 2013), between *Pdgfra*^*+/+*^ and *Pdgfra*^*PI3K/PI3K*^ E13.5 palatal mesenchyme samples by qPCR using primers spanning constitutively-expressed exons flanking the alternatively-spliced exon, confirming the trends observed in the rMATS analysis in four out of five biological replicates for each transcript (Figure 1C).

Next, differential gene expression was assessed in the above RNA-seq samples via DESeq2 (Love et al., 2014). Surprisingly, this analysis identified only 13 genes with significant differences in expression between the two genotypes (Table S4), none of which were differentially alternatively-spliced. Among these 13 genes, three (*Foxp2, Aldh1a2* and *Pdgfra*) have a corresponding mouse model with a craniofacial phenotype, with only *Pdgfra* models exhibiting a cleft secondary palate (Klinghoffer et al., 2002; Niederreither et al., 1999; Shu et al., 2005; Soriano, 1997). Notably, this finding indicates that the PI3K/Akt-mediated PDGFRα signaling axis primarily regulates gene expression through AS in the mid-gestation PS, at least in part through the regulation of Srsf3.

### Srsf3 *expression is enriched in the facial processes*

To assess where *Srsf3* is expressed during development, we performed whole mount RNA *in situ* hybridization from E8.5-E10.5. We saw an enrichment of transcripts in the head region at all timepoints (Figure 2A-2D), and particularly in the facial processes at E9.5-E10.5 (Figure 2B’ and 2C’), including the maxillary processes (MxP) from which the PS will extend. We then performed immunofluorescence analysis to examine expression of Srsf3 protein during development. From E8.5-E10.5, we observed predominantly nuclear Srsf3 expression in the pharyngeal arch and facial process mesenchyme as well as the overlying ectoderm (Figure 2E-2H). At E13.5, Srsf3 expression was detected along the anterior-posterior axis of the PS, with noticeably increased expression in the anterior PS (Figure 2I-2K).

**Figure 2.**
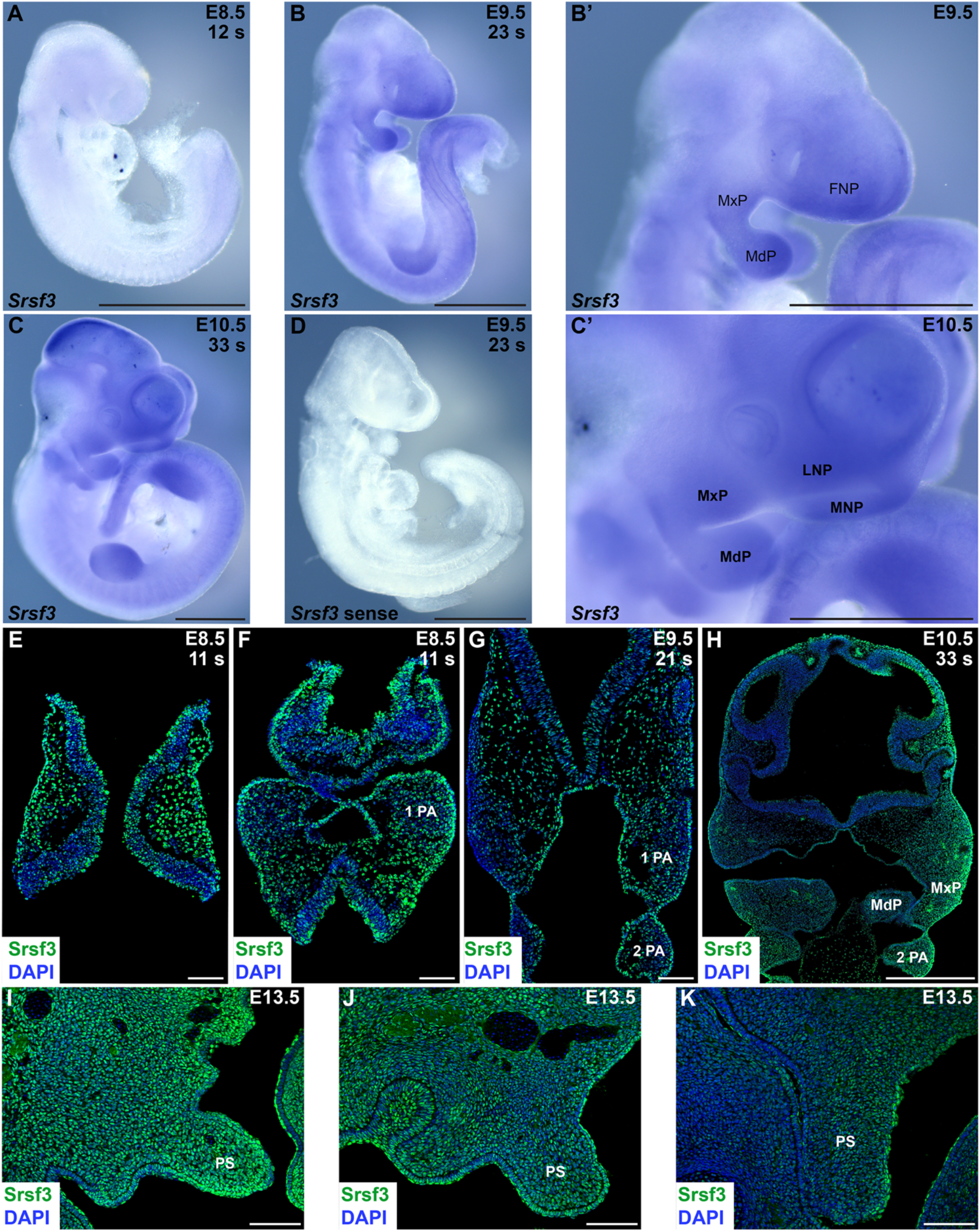
*Srsf3* expression is enriched in the facial processes. (A-C’) *Srsf3* expression as assessed by whole-mount *in situ* hybridization analysis with an antisense probe at E8.5 (A), E9.5 (B,B’) and E10.5 (C,C’). (D) No signal was detected with a control sense probe at E9.5. (E-K) Srsf3 expression (green) as assessed by immunofluorescence analysis on sections of the neural tube and pharyngeal arches at E8.5 (E,F) and E9.5 (G), the facial processes at E10.5 (H), and the anterior (I), middle (J) and posterior (K) PS at E13.5. Nuclei were stained with 4’6-diamidino-2-phenylindole (DAPI; blue). s, somite pairs; FNP, frontonasal prominence; MxP, maxillary process, MdP, mandibular process; 1 PA, first pharyngeal arch; 2 PA, second pharyngeal arch; PS, secondary palatal shelf. Bars, 1 mm (A-D), 100 μm (E-G, I-K), 500 μm (H).

### Phosphorylation of Srsf3 at Akt consensus sites in response to PDGF ligand stimulation leads to nuclear translocation

The full complement of Akt consensus sites (RxRxxS/T) (Alessi et al., 1996) found in the mouse Srsf3 RS domain are conserved from human through *Xenopus*, with five of the seven sites conserved in zebrafish (Figure 3A), demonstrating a high level of conservation among vertebrate species. To validate the PI3K-mediated phosphorylation of Srsf3 upon PDGF-AA ligand treatment detected in our previous mass spectrometry analysis (Fantauzzo and Soriano, 2014), we immunoprecipitated phosphorylated Akt substrates from nuclear and cytoplasmic fractions of immortalized MEPM (iMEPM) cells that were unstimulated or stimulated for 20 minutes with PDGF-AA ligand in the absence or presence of the PI3K inhibitor LY294002 (Vlahos et al., 1994) with an anti-Akt-phosphosubstrate antibody (Zhang et al., 2002) followed by Western blotting with an anti-Srsf3 antibody. This analysis revealed increased band intensities over baseline levels in response to PDGF-AA ligand treatment in the cytoplasmic fractions, indicative of increased phospho-Srsf3 levels, and decreased band intensities upon treatment with LY294002 (Figure 3B).

**Figure 3.**
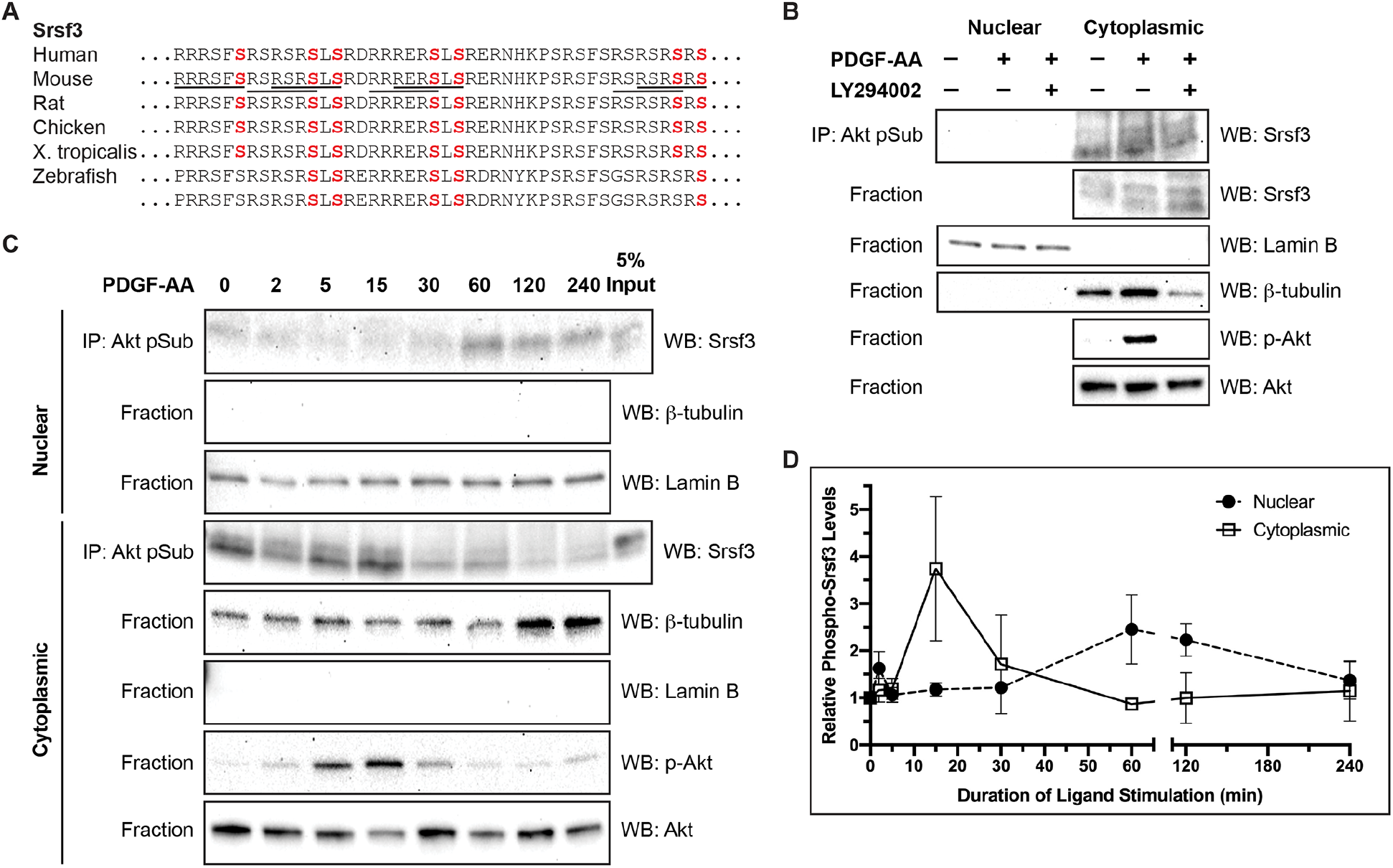
Phosphorylation of Srsf3 at Akt consensus sites in response to PDGF ligand stimulation leads to nuclear translocation. (A) Akt consensus sequence conservation in Srsf3 across vertebrate species. Full consensus sequences are underlined in mouse. Terminal serine residues in each Akt consensus sequence are highlighted in red. (B) Immunoprecipitation of Akt phosphorylation targets from nuclear and cytoplasmic fractions of iMEPM cells that were untreated, treated for 20 min with PDGF-AA ligand or pre-treated with the PI3K inhibitor LY294002 with an anti-Akt-phosphosubstrate antibody followed by Western blotting with an anti-Srsf3 antibody. Western blot analysis of fraction lysates confirmed the purity of the subcellular fractions as well as an increase in phospho-Akt levels upon PDGF-AA treatment. IP, immunoprecipitation; WB, Western blot. (C) Similar biochemical experiments as in (B), with a time course analysis of PDGF-AA ligand treatments from 2 to 240 minutes. (D) Line graph depicting quantification of band intensities from three independent experiments as in (C). Data are presented as mean ± SEM. See also Figure S1.

To investigate the effects of PI3K/Akt-mediated phosphorylation of the above sites on the subcellular localization of Srsf3, iMEPM cells were left unstimulated or stimulated with PDGF-AA ligand from two minutes to four hours. Immunoprecipitation of phosphorylated Akt substrates from the fractionated lysates using the anti-Akt phosphosubstrate antibody followed by Western blotting with the anti-Srsf3 antibody revealed that while phosphorylated Srsf3 was detected in both fractions at all timepoints, the cytoplasmic and nuclear fractions had the greatest amount of phosphorylated Srsf3 at 15 minutes (3.741 ± 1.527-fold induction over unstimulated levels) and 60 minutes (2.455 ± 0.7326-fold induction over unstimulated levels), respectively (Figure 3C and 3D). Significantly, the changes observed in phosphorylated Srsf3 levels in the nucleus over time in response to PDGF-AA ligand stimulation do not result from changes in total Srsf3 protein levels in this compartment following two hours of ligand stimulation (Figure S1A and S1B). These results suggest that phosphorylation of Srsf3 at Akt consensus sites downstream of PDGFRα signaling drive translocation of phosphorylated Srsf3 into the nucleus.

### *Ablation of* Srsf3 *in the neural crest lineage results in a midline facial clefting phenotype*

To characterize the role of Srsf3 during craniofacial development *in vivo*, we combined a *Srsf3* conditional allele (Jumaa et al., 1999) with the *Wnt1-Cre* transgene (Danielian et al., 1998) to ablate *Srsf3* in the NCC lineage. *Srsf3* itself is alternatively spliced to generate two transcripts. The major transcript excludes exon 4, while the minor transcript includes exon 4, which contains a premature termination codon and results in transcript degradation through nonsense-mediated decay (Figure S2A) (Jumaa and Nielsen, 1997). Upon Cre-mediated recombination, exons 2 and 3 in the conditional allele are deleted, resulting in a subsequent frameshift that generates a new start codon in exon 7 (Figure S2A). The ensuing 54-amino acid protein does not contain any of the domains present in the wild-type protein (Figure S2B). Analysis of *Srsf3* expression from RNA isolated from the facial processes and limb buds of E12.5 *Srsf3*^*fl/fl*^;*Wnt1-Cre*^*+/+*^ control and *Srsf3*^*fl/fl*^;*Wnt1-Cre*^*+/Tg*^ conditional knock-out (cKO) embryos confirmed efficient deletion of *Srsf3* exons 2 and 3 specifically in the facial processes of cKO embryos and exclusion of exon 4 in all samples tested (Figure S2C).

We crossed *Srsf3*^*fl/fl*^ males with *Srsf3*^*+/fl*^;*Wnt1-Cre*^*+/Tg*^ females and initially harvested progeny at birth. Genotyping revealed that cKO pups were recovered well below Mendelian frequency at this timepoint (2 cannibalized pups vs. 14 expected pups out of 54 total, p = 0.0003), with no live pups present at birth (Table S5). Additional harvests at embryonic timepoints indicated that the majority of cKO embryos died just past mid-gestation (Table S5). We first assessed the craniofacial phenotypes of cKO embryos by examining gross morphology during development. At E10.5 cKO embryos had hypoplastic facial processes, a widening of the space between the nasal pits, an enlarged forebrain and a reduced midbrain (Figure 4B-4B’’, Figure S3B) compared to their control littermates (Figure 4A-4A’’, Figure S3A). At E12.5 cKO embryos exhibited facial clefting, wherein the medial nasal processes had failed to fuse at the midline (Figure 4D’ and 4D’’). Moreover, the PS, tongue and midbrain were hypoplastic and the forebrain was misshapen (Figure 4D-4D’’, Figure S3D’-D’’’). Sectioning of cKO embryos at this timepoint revealed that the lateral ventricles had formed, though they were enlarged compared to those of control littermates, indicating that the anterior neural tube had successfully closed (Figure S3D-S3D’’’). By E14.5 the facial clefting phenotype in cKO embryos was more pronounced, with a striking cleft at the midline of the upper jaw and a subtler cleft in the mandible (Figure 4F’ and 4F’’). The tongue remained severely hypoplastic at this timepoint (Figure 4F’’) and continued forebrain overgrowth and exencephaly were observed (Figure 4F-4F’’). A subset of cKO embryos additionally exhibited facial subepidermal blebbing from E9.5-E12.5 (peaking at 43% at E12.5, n = 7), facial hemorrhaging from E10.5-E12.5 (peaking at 29% at E10.5, n = 31) and a wavy neural tube from E9.5-E11.5 (peaking at 24% at E11.5, n = 21). Importantly, the defects above were not due to a general delay in cKO embryo development nor global growth reduction, as structures such as the limb buds were indistinguishable from those of control embryos.

**Figure 4.**
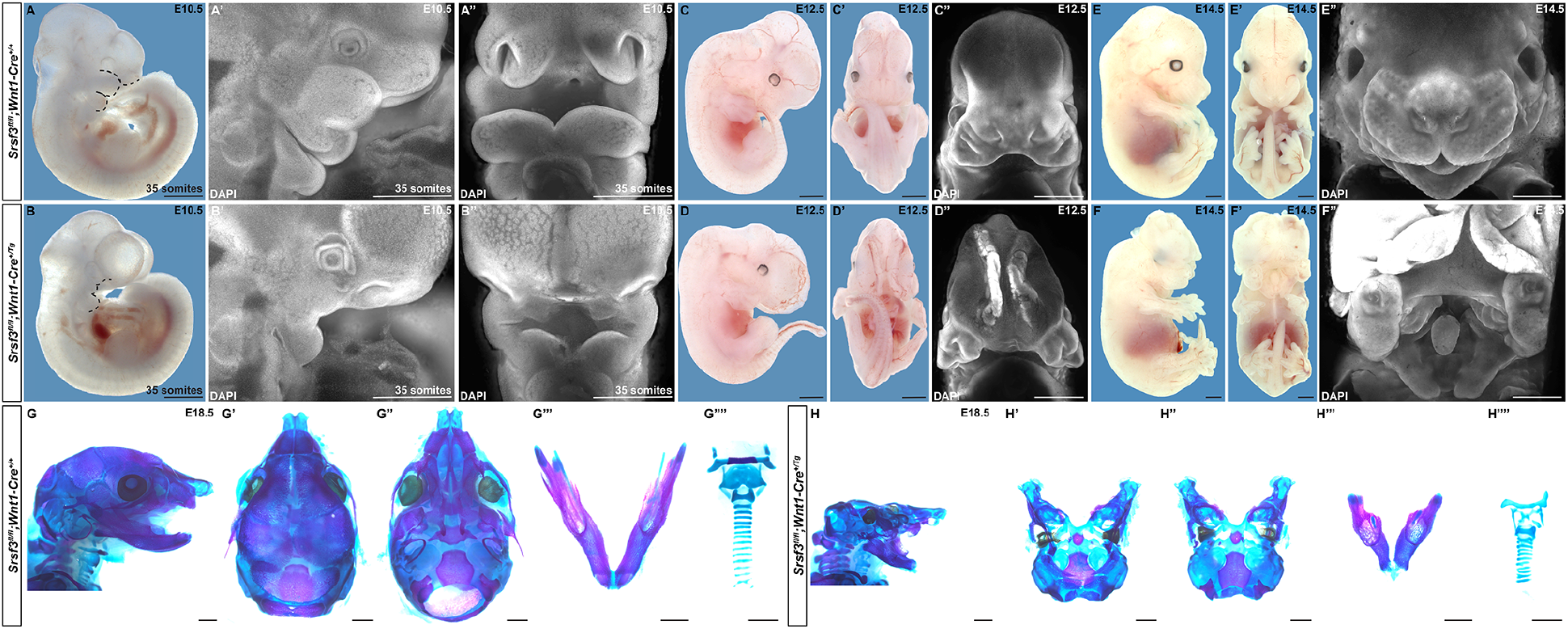
Ablation of *Srsf3* in the neural crest lineage results in a midline facial clefting phenotype. (A-F’’) Gross morphology of E10.5 (A-B’’), E12.5 (C-D’’) and E14.5 (E-F’’) *Srsf3*^*fl/fl*^;*Wnt1-Cre*^*+/+*^ control embryos (top) and *Srsf3*^*fl/fl*^;*Wnt1-Cre*^*+/Tg*^ cKO embryos (bottom) as viewed either laterally before (A-F) and after DAPI staining (A’,B’) or frontally before (C’-F’) and after DAPI staining (A’’-F’’). Dotted black lines (A,B) outline nasal pits and first pharyngeal arch. (G-H’’’’) Lateral (G,H), dorsal (G’,H’) and ventral (G’’,H’’) views of craniofacial skeletal preparations as well as dissected mandibles (G’’’,H’’’) and dissected hyoid bones and thyroid, cricoid and tracheal cartilages (G’’’’,H’’’’) generated from E18.5 control embryos (left) and cKO embryos (right). Bars, 1 mm. See also Figures S2 and S3.

We next evaluated the craniofacial phenotypes of three cKO embryos that survived to approximately E18.5 by generating skeletal preparations. In general, cKO embryos exhibited a smaller head than their littermate controls (Figure 4G and 4H). The anterior part of the cKO face was clefted and those bones and cartilages tended to be hypoplastic. These elements included the premaxilla, frontal process of premaxilla, maxilla, frontal process of maxilla and zygomatic process of maxilla bones as well as the nasal cartilage. The nasal and frontal bones were missing entirely. The middle of the face had elements that were more severely affected and in several cases absent. These hypoplastic structures included the jugal, zygomatic process of squamosal, squamosal, parietal, basisphenoid, pterygoid and retrotympanic process bones as well as the cartilages of the middle ear. The palatal process of the palatine, palatine, ala temporalis, lamina obturans and ectotympanic bones were absent from the middle of the face as was the processus styloidius cartilage. The palatal process of the maxilla was absent in two of three cKO embryos and hypoplastic in the remaining embryo. The posterior part of the skull derived from the paraxial mesoderm was relatively unaffected, though the interparietal and supraoccipital bones were clefted and hypoplastic (Figure 4H’ and 4H’’). The mandible, coronoid process and Meckel’s cartilage of cKO embryos were hypoplastic, with clefting of the mandible observed in one embryo (Figure 4H’’’). The hyoid was present but not ossified in two cKO embryos and severely hypoplastic in the third embryo, while the lesser and greater horns of the hyoid were hypoplastic (Figure 4H’’’’). Finally, the thyroid and cricoid cartilages were hypoplastic and the tracheal cartilage rings were misshapen and occasionally fused (Figure 4H’’’’).

### *Cranial neural crest cell specification and survival are defective in* Srsf3 *conditional knock-out embryos*

We next introduced the *ROSA26*^*mTmG*^ double-fluorescent Cre reporter allele (Muzumdar et al., 2007) through the male germline in the above crosses to examine NCC specification and migration. At E8.0, whereas *Srsf3*^*+/fl*^;*Wnt1-Cre*^*+/Tg*^;*ROSA26*^*+/mTmG*^ control embryos had bright, GFP-positive cells in the cranial neural folds (Figure 5A and 5A’), *Srsf3*^*fl/fl*^;*Wnt1-Cre*^*+/Tg*^;*ROSA26*^*+/mTmG*^ cKO littermates had noticeably fewer GFP-positive cells (Figure 5B and 5B’), indicative of a NCC specification defect. cKO embryos had fewer GFP-positive cells in the frontonasal prominence and first pharyngeal arch derivatives at E9.5, as well as a more diffuse stream of NCCs entering pharyngeal arch 2 (Figure 5D and 5D’) compared to control embryos (Figure 5C and 5C’). At E10.5, whereas control embryos had intense, GFP-positive cells throughout the facial processes and clearly-delineated NCC streams entering pharyngeal arches 3 and 4 (Figure 5E’), cKO littermates had very few GFP-positive cells in the facial processes and no obvious NCC streams entering the pharyngeal arches (Figure 5F’), indicating a defect in NCC-derived cell survival between E9.5-E10.5. GFP-positive cells were detected in the brain and dorsal root ganglia of cKO embryos (Figure 5D’ and 5F’), though with decreased intensity compared to control littermates (Figure 5C’ and 5E’).

**Figure 5.**
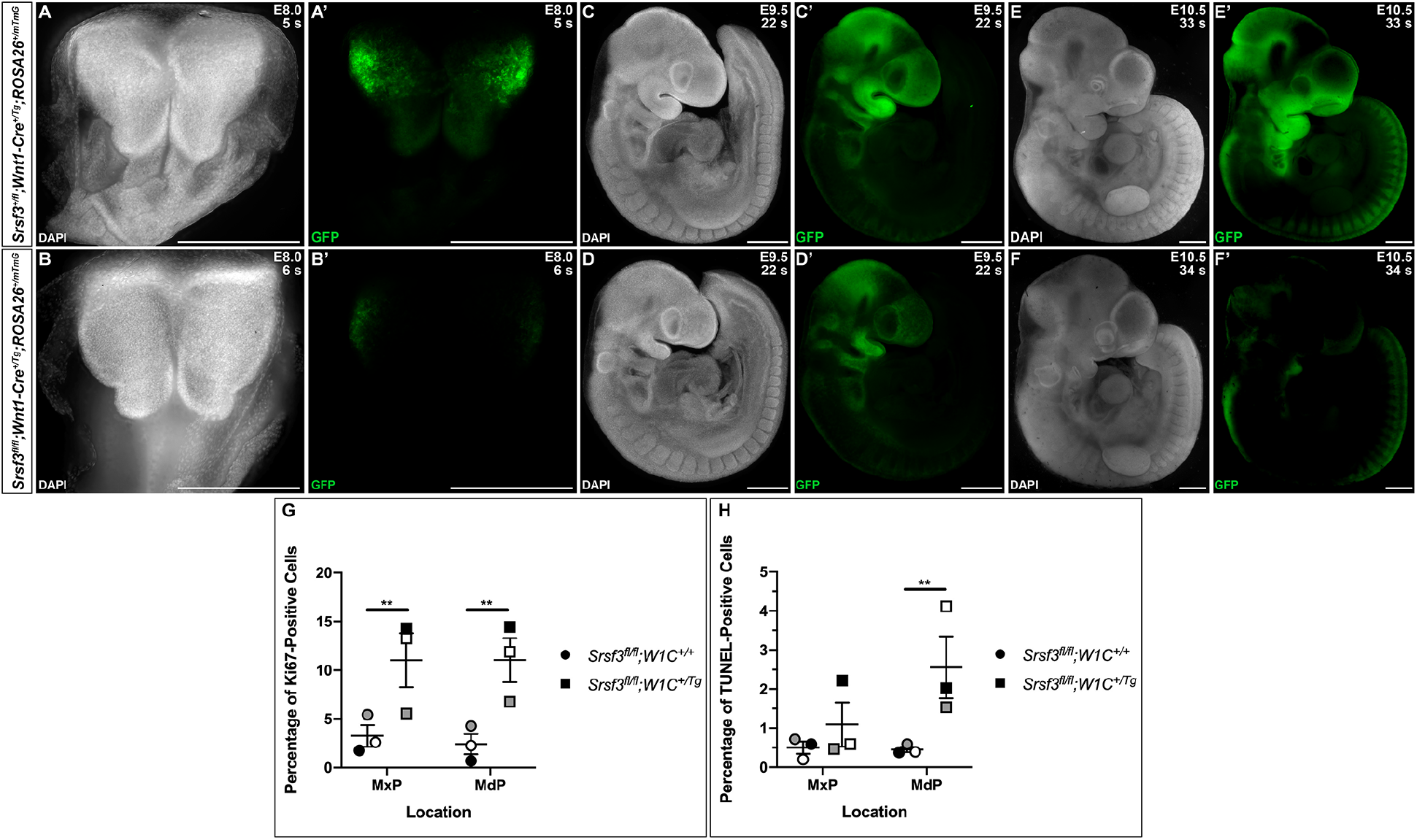
Cranial neural crest cell specification and survival are defective in *Srsf3* conditional knock-out embryos. (A-F’) Dorsal (A-B’) and lateral (C-F’) whole-mount fluorescence images of DAPI (A-F) and GFP (A’-F’) expression in *Srsf3*^*+/fl*^;*Wnt1-Cre*^*+/Tg*^;*ROSA26*^*+/mTmG*^ control embryos (top) and *Srsf3*^*fl/fl*^;*Wnt1-Cre*^*+/Tg*^;*ROSA26*^*+/mTmG*^ cKO embryos (bottom) at E8.0 (A-B’), E9.5 (C-D’) and E10.5 (E-F’). s, somite pairs. Bars, 500 μm. (G,H) Scatter dot plots depicting the average percentage of Ki67-positive (G) and TUNEL-positive (H) cells per embryo in the maxillary and mandibular processes at E10.5. Data are presented as mean ± SEM. **, p < 0.01. Shades correspond to independent experiments across three biological replicates of control and cKO samples. MxP, maxillary process; MdP, mandibular process.

We next assessed both cell proliferation and cell death in the maxillary and mandibular processes of *Srsf3*^*fl/fl*^;*Wnt1-Cre*^*+/+*^ control versus *Srsf3*^*fl/fl*^;*Wnt1-Cre*^*+/Tg*^ cKO embryos at E10.5 via Ki67 immunofluorescence analysis and terminal deoxynucleotidyl transferase-mediated dUTP nick end labeling (TUNEL), respectively. cKO embryos had statistically significant increases in the percentage of Ki67-positive cells compared to control embryos in both the maxillary (p = 0.004) and mandibular processes (p = 0.004) (Figure 5G). Similarly, there was a trend for the percentage of TUNEL-positive cells to be higher in the MxP of cKO embryos compared to control embryos, with a statistically significant increase in this value in the mandibular processes (p = 0.006) (Figure 5H).

### Srsf3 regulates the alternative RNA splicing of transcripts encoding protein kinases in the facial process mesenchyme

Given the severity of the facial phenotype in cKO embryos, we next examined whether AS remained intact in this setting. Analysis of *Fgfr2* expression from RNA isolated from the facial processes of E12.5 control and cKO embryos revealed that both epithelia- and mesenchyme-specific splice isoforms were observed in each genotype (Figure S2D), indicating that Srsf3 likely regulates the AS of a specific subset of transcripts in the facial mesenchyme. To identify those transcripts, we harvested and sequenced MxP mesenchyme RNA from three biological replicates of E11.5 control versus cKO embryos (Table S1). Consistent with the qPCR results above (Figure S2C), reads from control embryos at the *Srsf3* locus included exons 2 and 3, while reads from cKO embryos often skipped these exons. Further, there were very few reads including *Srsf3* exon 4 in both control and cKO samples (Figure S4).

Differential gene expression between control and cKO embryos was assessed via DESeq2 as above. This analysis identified 423 genes with significant differences in expression between the two genotypes (Table S6; Figure 6A). As expected, *Srsf3* itself was not differentially expressed between control and cKO samples (adjusted p-value = 0.94). A GO analysis of these 423 genes using the KEGG 2019 Mouse, GO Biological Process 2018 and GO Molecular Function 2018 libraries of Enrichr indicated that the top terms related to focal adhesions, the extracellular matrix and cellular signaling (Figure 6B). We examined the differential expression of representative genes from these terms with associated mouse craniofacial phenotypes, including *Pdgfc, Col2a1* and *Kdr* (Aszódi et al., 2001; Ding et al., 2004; Sandell et al., 2011), between control and cKO E11.5 MxP mesenchyme samples by qRT-PCR. This analysis confirmed the trends observed in the DESeq2 analysis for all three genes, with significantly increased levels of *Col2a1* in the cKO samples compared to the control samples (p = 0.04) (Figure 6C). Among the 423 differentially-expressed genes, 65 have mouse models with craniofacial phenotypes, 21 of which exhibit a cleft secondary palate (Table S7). A GO analysis of these 65 genes using the WikiPathways 2019 library of Enrichr revealed that the second most significant term was neural crest differentiation (p = 1.9 × 10^−5^). We examined the differential expression of three genes from this term, *Foxd3, Col2a1, Zic5*, and one additional gene involved in mouse craniofacial development, *Pou3f4* (Aszódi et al., 2001; Inoue et al., 2004; Phippard et al., 1999; Teng et al., 2008), between control and cKO E11.5 MxP mesenchyme samples by qRT-PCR, confirming a significant increase in *Zic5* and *Pou3f4* expression in cKO samples compared to the control samples (p = 0.04) (Figure 6C).

**Figure 6.**
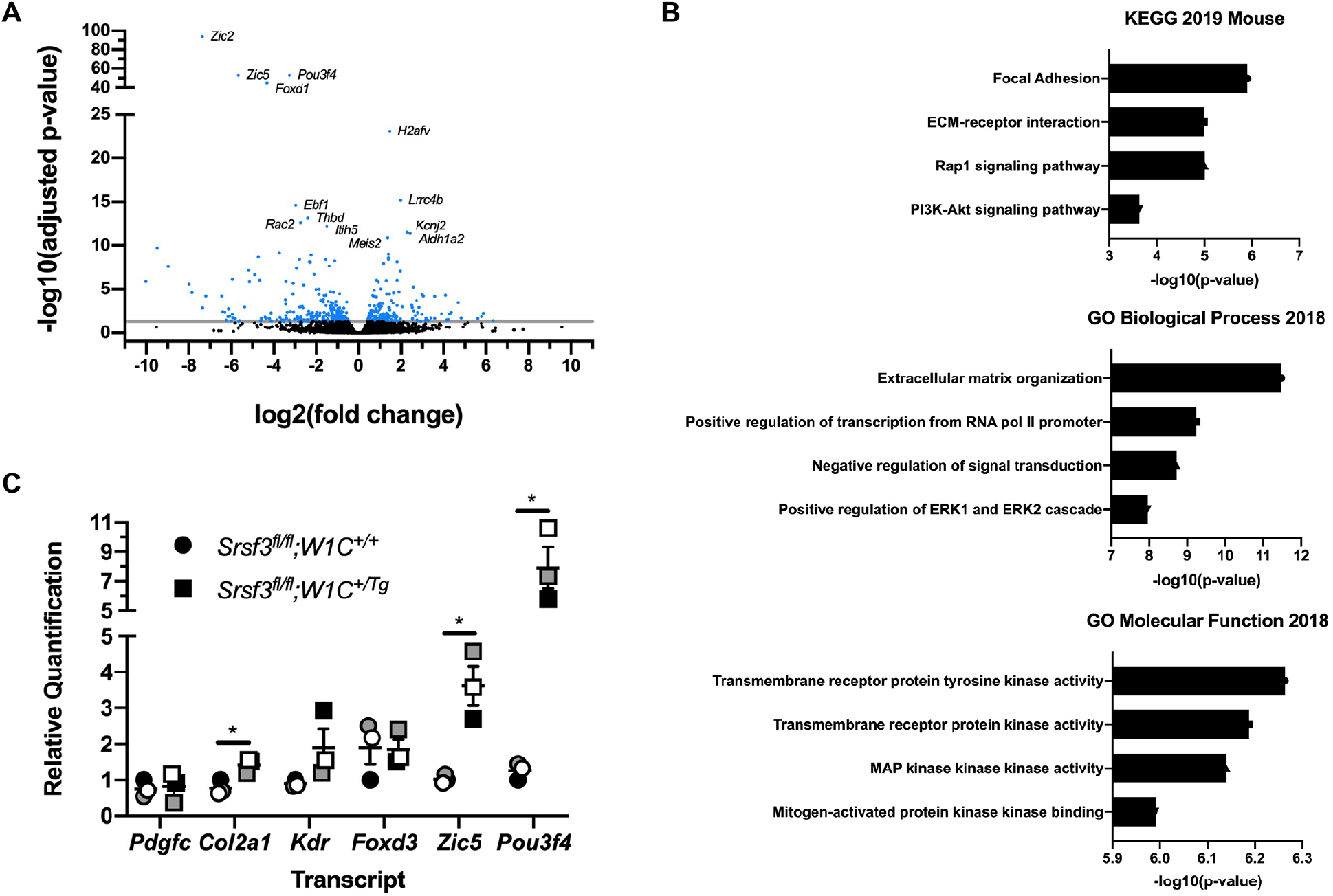
Differential transcript expression in *Srsf3* conditional knock-out embryos points to changes in focal adhesions, the extracellular matrix and cellular signaling. (A) Volcano plot depicting the −log10(adjusted p-value) versus log2(fold change) detected by DESeq2 analysis of RNA-seq data from E11.5 *Srsf3*^*fl/fl*^;*Wnt1-Cre*^*+/+*^ control versus *Srsf3*^*fl/fl*^;*Wnt1-Cre*^*+/Tg*^ cKO MxP mesenchyme. Dots in blue have an adjusted p-value ≤0.05. (B) Bar graphs depicting GO terms with associated −log10(p-value) detected by a GO analysis of differentially-expressed transcripts represented in (A) using various Enrichr libraries. (C) Scatter dot plot depicting qRT-PCR values for six differentially-expressed transcripts between E11.5 control versus cKO MxP mesenchyme. Data are presented as mean ± SEM. *, p < 0.05. Shades correspond to independent experiments across three biological replicates of control and cKO samples. See also Figure S4.

rMATS was again used to detect AS events in this same dataset, revealing 1,392 events that were significantly different between genotypes across the five major classes, with the majority of events (65.7%) involving SE (Table S8; Figure 7A). Only 30 transcripts (2.9% of all rMATs events) were detected in both the DESeq2 and rMATS analyses (Table S6, bold). We used RBPmap as above to examine annotated Srsf3 motifs in the 915 SE plus 250 bp flanking each end, revealing Srsf3 motifs in 895 (97.8%) of these ROIs (Table S8). The density of these motifs was greater in significant (5.1 motifs per ROI) versus non-significant (4.5 motifs per ROI) SE events in our dataset (p = 1.3 × 10^−4^). Again, Srsf3 motifs were enriched flanking the SE in the significant SE events (p-value < 0.0001) (Figure 7B). Among the 728 genes represented in the SE class that had one or more Srsf3 motifs in the ROI, 45 have a corresponding mouse model with a craniofacial phenotype, 14 of which have a cleft or arched secondary palate (Table S9). A GO analysis of the genes represented in the SE class using the GO Biological Process 2018, GO Molecular Function 2018 and InterPro Domains 2019 libraries of Enrichr revealed that the top terms were phosphorylation (p = 1.3 × 10^−4^), protein serine/threonine kinase activity (p = 5.7 × 10^−5^) and protein kinase domain (p = 6.4 × 10^−4^), respectively. We examined the differential AS of *Melk, Limk2*, *Dmpk* and *Prkd2*, each of which was included in these three GO terms, between control and cKO E11.5 MxP mesenchyme samples by qPCR using primers spanning constitutively-expressed exons flanking the alternatively-spliced exon. These analyses confirmed the trends observed in the rMATS analysis in two out of three biological replicates for *Melk* and all three biological replicates for *Limk2, Dmpk* and *Prkd2*, the last of which had a significant difference in percent spliced between genotypes (p = 0.02) (Figure 7C).

**Figure 7.**
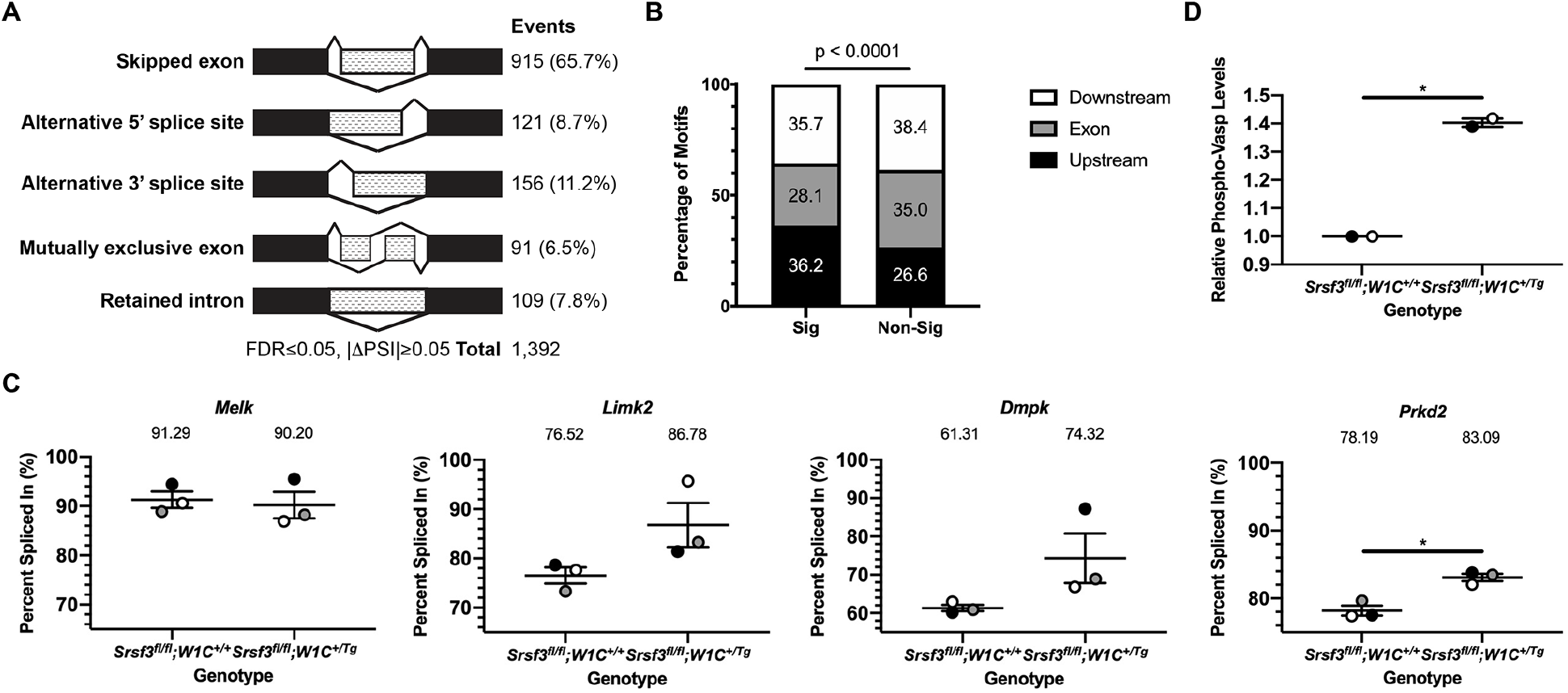
Srsf3 regulates the alternative RNA splicing of transcripts encoding protein kinases in the facial process mesenchyme. (A) Summary of differential alternative splicing events detected by rMATS analysis of RNA-seq data from E11.5 *Srsf3*^*fl/fl*^;*Wnt1-Cre*^*+/+*^ control versus *Srsf3*^*fl/fl*^;*Wnt1-Cre*^*+/Tg*^ cKO MxP mesenchyme. FDR, false discovery rate; ΔPSI, difference in percent spliced in. (B) Bar graph depicting the percentage of Srsf3 motifs detected by RBPmap within or flanking exons between significant (Sig) and non-significant (Non-Sig) skipped exon events. (C) Scatter dot plots depicting percent spliced in for four differentially alternatively-spliced transcripts between E11.5 control versus cKO MxP mesenchyme as assessed by qPCR. Data are presented as mean ± SEM. *, p < 0.05. Shades correspond to independent experiments across three biological replicates of control and cKO samples. (D) Scatter dot plot depicting relative phospho-Vasp levels between E11.5 control versus cKO MxP mesenchyme. Data are presented as mean ± SEM. *, p < 0.05. Shades correspond to independent experiments, each consisting of pooled lysates from three embryos per genotype.

We next employed SPLINTER (Low, 2020) to predict outcomes stemming from the AS of the SE events (Table S8), with a particular focus on the 27 events for transcripts encoding 21 protein serine/threonine kinases. Nonsense-mediated decay was predicted in 12 cases, potentially leading to the downregulation of expression for 9 genes in control embryos and 3 genes in cKO embryos. A truncated protein was predicted in 7 cases, applying to 2 proteins in control embryos and 5 proteins in cKO embryos. Here, the protein kinase domain in particular would be affected in 2 proteins from each genotype. Among these protein serine/threonine kinases, Wnk1, Map3k12, Prkaa1 and Irak1 are demonstrated Akt substrates (Chen et al., 2002; Heathcote et al., 2016; Vitari et al., 2004; Wu et al., 2015), as are several of their phosphorylation substrates. Of particular interest, we detected increased phosphorylation of the protein serine/threonine kinase Prkd2 and its substrate Vasp upon PDGF-AA ligand treatment of primary MEPM cells in our previous phosphoproteomic screen (Fantauzzo and Soriano, 2014). Also, *Vasp* was found to be differentially alternatively spliced in our E13.5 PS mesenchyme rMATS analysis, resulting in a loss of amino acids containing the Akt consensus motif and the final Prkd2 consensus motif more often in *Pdgfra*^*PI3K/PI3K*^ samples (Table S2), indicating further regulation of *Vasp* downstream of PI3K/Akt-mediated PDGFRα signaling. Consistent with the SPLINTER prediction that AS of *Prkd2* would result in a truncated protein in control embryos affecting amino acids that fall within the protein kinase domain and include the proton acceptor, phosphorylation of Vasp was increased in pooled *Srsf3* cKO E11.5 MxP mesenchyme lysates compared to those of control embryos (p = 0.02) (Figure 7D). These results indicate that Srsf3 serves to negatively regulate PDGFRα signaling through the AS of protein serine/threonine kinases.

## Discussion

Despite the breadth of studies on PDGF signaling, the intracellular effectors that bridge the receptor-bound signaling molecules and gene expression changes downstream of PDGFRα activation remain largely unknown. Only two studies have shed light on the transcriptional changes downstream of this signaling pathway in the context of craniofacial development, both using primary MEPM cells. PDGFRα signaling was shown to specifically upregulate the expression of 41 genes bound by the SRF transcription factor, a subset of which were enriched for MRTF cofactor binding sites and associated with the cytoskeleton (Vasudevan and Soriano, 2014). Further, PDGFRα signaling was demonstrated to principally depend on PI3K activity and ultimately promote osteoblast differentiation through the upregulation of 24 transcripts associated with cell differentiation (Vasudevan et al., 2015). Here, we revealed that the PI3K/Akt-mediated PDGFRα signaling axis primarily regulates gene expression through AS *in vivo* in the mid-gestation PS and discovered 523 transcripts that are differentially alternatively spliced when this pathway is disrupted. This study thus highlights a novel role for PDGFRα signaling in regulating RNA processing and identifies the phosphorylation of RBPs downstream of this pathway as a mechanism contributing to this response.

Transcripts encoding each of the 11 SR proteins expressed in mouse were detected in both RNA-seq analyses presented here. In the E13.5 PS mesenchyme, *Srsf1*, *Srsf2* and *Srsf6* had average normalized read counts in the control samples on the same order of magnitude as *Srsf3*. Further, each of the murine SR proteins contains at least one Akt consensus sequence and as many as 22 in the case of Srsf4. In addition to human SRSF3 (Bavelloni et al., 2014), human SRSF1 and SRSF7 have previously been shown to be phosphorylated by Akt in *in vitro* kinase assays (Blaustein et al., 2005). It is therefore interesting that Srsf3 was the only SR protein detected in our previous mass spectrometry screen as an Akt phosphorylation target downstream of PI3K-mediated PDGFRα signaling in the palatal mesenchyme (Fantauzzo and Soriano, 2014). Further confirming this finding, the severe facial clefting phenotype of *Srsf3* cKO embryos indicates that other SR proteins expressed in the facial mesenchyme are unable to compensate for the loss of Srsf3 in this setting. This idea is consistent with previous results demonstrating that two human SR proteins, SRSF3 and SRSF4, have distinct consensus binding motifs and mostly unique target transcripts in the P19 mouse embryonic carcinoma cell line (Änkö et al., 2012). Further, none of the transcripts encoding SR proteins were differentially expressed in either DESeq2 analysis here nor differentially alternatively spliced in the *Pdgfra*^*+/+*^ versus *Pdgfra*^*PI3K/PI3K*^ rMATS analysis. Only *Srsf3* was found to be differentially alternatively spliced in the *Srsf3*^*fl/fl*^;*Wnt1-Cre*^*+/+*^ versus *Srsf3*^*fl/fl*^;*Wnt1-Cre*^*+/Tg*^ rMATS analysis. These results demonstrate that, unlike in P19 cells (Änkö et al., 2012), Srsf3 is not a master regulator of SR protein-encoding transcript splicing in the facial mesenchyme.

Interestingly, the Srsf3 Western blot bands present in the absence of PDGF-AA ligand stimulation following Akt phosphosubstrate immunoprecipitation further confirm that the majority of Akt phosphorylation targets in MEPM cells are phosphorylated at baseline levels (Fantauzzo and Soriano, 2014). This finding suggests that input(s) other than PDGF-AA-stimulated PDGFRα signaling contributes to Srsf3 phosphorylation at Akt consensus sites. SR proteins have been shown to be phosphorylated in response to multiple treatments, such as downstream of hepatocyte, keratinocyte and epidermal growth factor stimulation and all-*trans* retinoic acid treatment (Bavelloni et al., 2014; Blaustein et al., 2005; Zhou et al., 2012). As these growth factors, metabolites and/or their receptors have demonstrated roles in regulating the activity of NCCs and the development of their derivatives in the mouse (Fantauzzo and Soriano, 2015; Williams and Bohnsack, 2019), it is likely that one or more of these pathways, or an as yet undetermined signaling pathway, contributes to Srsf3 phosphorylation in the palatal mesenchyme. While we have examined the effect of Srsf3 phosphorylation in response to PDGF-AA ligand stimulation on its subcellular localization, future studies will determine whether these same post-translational modifications downstream of PI3K/Akt-mediated PDGFRα signaling also affect Srsf3 RNA binding and/or sequence specificity.

Our study has revealed a novel requirement for an SR protein in NCCs and the facial mesenchyme. Upon ablation of *Srsf3* in the NCC lineage, a reduced number of cranial NCCs are initially specified, which are able to migrate into the facial processes, albeit to less of an extent than in control embryos. By E10.5 cKO embryos exhibit increased cell death in the facial processes which is partially offset by increased cell proliferation, thereby allowing derivatives of these processes to form in embryos that survive past mid-gestation. In accordance with Srsf3 acting as an effector downstream of PDGFRα signaling, *Srsf3* cKO and *Pdgfra* mutant mouse models have a number of overlapping phenotypes, including midline facial clefting, PS hypoplasia, facial subepidermal blebbing, facial hemorrhaging and a wavy neural tube (Fantauzzo and Soriano, 2014; Klinghoffer et al., 2002; Soriano, 1997). However, the craniofacial skeletal phenotypes in *Srsf3* cKO embryos are more severe than those found upon ablation of *Pdgfra* in the NCC lineage (He and Soriano, 2013; Tallquist and Soriano, 2003), again indicating that Srsf3 acts to mediate more than PDGFRα signaling in this setting. One of the most striking phenotypes in *Srsf3* cKO embryos, and one not found in *Pdgfra* mutant models, are the brain malformations. Perhaps consistent with these defects, out of 109 *Srsf3*^*+/fl*^;*Wnt1-Cre*^*+/Tg*^ females in our colony, 16% died prematurely with an average age at death of 46 days. These females were often found with foam and/or blood at the mouth, indicative of a neurological issue. The enlarged lateral ventricle phenotype in cKO embryos is reminiscent of the phenotype observed in *Foxc1* mutant mice, which stems from defects in development of the forebrain meninges and reduced retinoic acid secretion from this structure (Siegenthaler et al., 2009). Interestingly, we found that *Foxc1* and *Aldh1a2* were both differentially expressed between control and *Srsf3* cKO embryo MxP mesenchyme samples. Together, these results suggest that Srsf3 may be required for meninges development, a hypothesis that will be tested in future studies.

Comparing the two RNA-seq experiments performed here, there were 38 AS events commonly detected in both rMATS analyses. Among these, only 12 (32%) were differentially alternatively spliced in the same direction (included or excluded) in *Pdgfra*^*PI3K/PI3K*^ and *Srsf3* cKO embryos compared to their respective control embryos. For the DESeq2 analyses, only two transcripts were commonly detected, *Aldh1a2* and *Rragd*. The relatively low extent of overlap between identified transcripts is not surprising given that these RNA-seq experiments were performed in different, but related, tissues across a 48-hour timeframe. In fact, AS changes in the developing murine face from E10.5-E12.5 are more significant across age than across facial prominence location (Hooper et al., 2020).

Further, there was little to no overlap between the transcripts detected in the rMATS and DESeq2 analyses within either RNA-seq experiment, indicating that loss of Srsf3 or a decrease in its activity does not globally affect the stability of its splicing targets in the facial mesenchyme. The majority of transcripts identified in the DESeq2 analyses are likely regulated by Srsf3 through mechanisms other than AS, such as transcription, export, translation and degradation (Howard and Sanford, 2015). Notably, among the SE events with one or more Srsf3 motifs in the ROI detected in our E13.5 PS mesenchyme rMATS analysis, significantly more events had a positive (260), as opposed to negative (121), PSI in which the differentially alternatively-spliced exon was included more often in the *Pdgfra*^*+/+*^ control samples. As SR proteins promote exon inclusion (Fu and Ares, 2014; Licatalosi and Darnell, 2010), this finding supports our hypothesis that the ability of Srsf3 to regulate AS is partially disrupted in *Pdgfra*^*PI3K/PI3K*^ embryos. Going forward, it will be critical to identify transcripts that are directly bound by Srsf3 in the facial mesenchyme through eCLIP analyses. Of interest, of the genes represented by the differentially alternatively-spliced transcripts identified in our E11.5 MxP mesenchyme rMATS analysis, transcripts from 63.0%, 48.8% and 13.5% were shown to be bound by human SRSF3 in mouse embryonic stem cells, HeLa cells and P19 cells, respectively (Änkö et al., 2012; Ratnadiwakara et al., 2018; Xiao et al., 2016).

Taken together, our findings significantly expand our understanding of the molecular mechanisms by which PDGFRα signaling controls gene expression and identify Srsf3 as a novel intracellular effector of this pathway during mammalian midface development. Future studies will seek to establish an AS “code” in the facial mesenchyme to address how RBPs might function in a cooperative or antagonistic fashion in this context and determine how the activity of these proteins is affected by post-translational modification downstream of receptor tyrosine kinase activity.

## Acknowledgments

We thank Damian Garno and Robert A. Long for technical assistance. RNA-seq experiments were performed at the Genomics and Microarray Core at the University of Colorado Anschutz Medical Campus. *Srsf3*^*fl*^ mice were a gift from Dr. Nicholas J. Webster at the University of California San Diego. We are grateful to Drs. David Clouthier and Julie Siegenthaler at the University of Colorado Anschutz Medical Campus for assistance with analysis of *Srsf3*^*fl/fl*^;*Wnt1-Cre*^*+/Tg*^ phenotypes. We thank members of the Fantauzzo laboratory, Dr. Matthew Taliaferro and Dr. David Clouthier for their critical comments on the manuscript. This work was supported by National Institute of Dental and Craniofacial Research grants R01DE027689 (to K.A.F.), K02DE028572 (to K.A.F) and F31DE029364 (to B.J.C.D.), and a RNA Bioscience Initiative Graduate Scholar Award (to B.J.C.D.) and RNA-seq grant (to K.A.F.).

## Author Contributions

Conceptualization, K.A.F.; Methodology, B.J.C.D., E.D.L., and K.A.F.; Investigation: B.J.C.D., J.M., and K.A.F.; Formal Analysis: B.J.C.D., E.D.L., R.F., and K.A.F.; Visualization: B.J.C.D. and K.A.F.; Writing – Original Draft: K.A.F.; Writing – Reviewing and Editing: B.J.C.D., E.D.L., R.F., J.M. and K.A.F.; Funding Acquisition: B.J.C.D. and K.A.F.; Supervision: K.A.F.

## Supplemental Figure Legends

**Figure S1, related to Figure 3.** Total Srsf3 protein levels in the nucleus do not change in response to PDGF-AA ligand treatment. (A) Western blot analysis of total Srsf3 levels in nuclear fractions of iMEPM cells that were untreated or treated with PDGF-AA ligand in a time course analysis from 2 to 240 minutes. WB, Western blot. (B) Line graph depicting quantification of band intensities from three independent experiments as in (A). Data are presented as mean ± SEM. *, p < 0.05.

**Figure S2, related to Figure 4.** Efficient deletion of *Srsf3* exons 2 and 3 and maintenance of alternative RNA splicing in *Srsf3* conditional knock-out embryos. (A) Conditional, floxed *Srsf3* locus before (top) and after (bottom) Cre exposure. Boxes represent exons. Colors correspond to protein domains in (B). Black dots indicate sites of forward (F) and reverse (R) primers used in (C). Triangles represent loxP sites. Dotted line surrounds exon 4, which contains a premature termination codon and is excluded from the major transcript. (B) Srsf3 wild-type (top) and cKO (bottom) proteins. Amino acid residues at the boundaries of the RNA recognition motif (RRM) and arginine/serine-rich (RS) domain are indicated. The epitope for the antibody used in Figure 2, Figure 3 and Figure S1 is indicated. (C) Representative qPCR product gel depicting *Srsf3* transcripts in the facial processes (left) and limb buds (right) of E12.5 *Srsf3*^*fl/fl*^;*Wnt1-Cre*^*+/+*^ control versus *Srsf3*^*fl/fl*^;*Wnt1-Cre*^*+/Tg*^ cKO embryos. (D) Representative qPCR product gel depicting *Fgfr2* and control *B2m* transcripts in the facial processes of E12.5 control versus cKO embryos.

**Figure S3, related to Figure 4**. Craniofacial morphological defects in *Srsf3* conditional knock-out embryos. (A-D’’’) Hematoxylin and eosin-stained coronal sections of E10.5 (A,B) and E12.5 (C-D’’’) *Srsf3*^*fl/fl*^;*Wnt1-Cre*^*+/+*^ control embryos (top) and *Srsf3*^*fl/fl*^;*Wnt1-Cre*^*+/Tg*^ cKO embryos (bottom). Sections in C-D’’’ move from anterior to posterior craniofacial structures. LNP, lateral nasal process; MNP, medial nasal process; MxP, maxillary process; MdP, mandibular process, LV, lateral ventricle, NS, nasal septum, TV, third ventricle, PS, secondary palatal shelf, T, tongue. Bars, 1 mm.

**Figure S4, related to Figure 6.***Srsf3* conditional knock-out RNA-seq reads skip exons 2 and 3. IGV snapshots of *Srsf3* transcripts from RNA-seq analysis of three biological replicates each of E11.5 *Srsf3*^*fl/fl*^;*Wnt1-Cre*^*+/+*^ control versus *Srsf3*^*fl/fl*^;*Wnt1-Cre*^*+/Tg*^ cKO MxP mesenchyme.

## STAR Methods

### LEAD CONTACT AND MATERIALS AVAILABILITY

Further information and requests for resources and reagents should be directed to and will be fulfilled by the Lead Contact, Katherine A. Fantauzzo. The *Srsf3 in situ* hybridization probe plasmid generated in this study will be made available on request with reasonable compensation by requestor for its processing and shipping.

### EXPERIMENTAL MODEL AND SUBJECT DETAILS

#### Mouse strains and husbandry

All animal experimentation was approved by the Institutional Animal Care and Use Committee of the University of Colorado Anschutz Medical Campus. *Pdgfra*^*tm5Sor*^ mice (Klinghoffer et al., 2002), referred to in the text as *Pdgfra*^*PI3K*^ mice; *Srsf3*^*tm1Pjln*^ mice (Jumaa et al., 1999), referred to in the text as *Srsf3*^*fl*^ mice; *H2afv*^*Tg(Wnt1-cre)11Rth*^ mice (Danielian et al., 1998), referred to in the text as *Wnt1-Cre*^*Tg*^ mice; and *Gt(ROSA)26Sor*^*tm4(ACTB-tdTomato,-EGFP)Luo*^ mice (Muzumdar et al., 2007), referred to in the text as *ROSA26*^*mTmG*^ mice, were maintained on a 129S4 coisogenic genetic background and housed at a sub-thermoneutral temperature of 21-23°C. Embryos were obtained from intercrosses of *Srsf3*^*fl/fl*^ males with *Srsf3*^*+/fl*^;*Wnt1-Cre*^*+/Tg*^ females or intercrosses of *Srsf3*^*fl/fl*^;*ROSA26*^*+/mTmG*^ males with *Srsf3*^*+/fl*^;*Wnt1-Cre*^*+/Tg*^ females. Mice were euthanized by inhalation of carbon dioxide from compressed gas. Cervical dislocation was used as a secondary verification of death. Both male and female embryos were analyzed in this study and no differences in phenotype were detected between these two groups. Developmental stages are described in the Results section for individual experiments. Control and experimental embryos were harvested from the same litter and embryos were age-matched to the greatest extent possible by somite counting and/or digit volume measurement. Statistical analyses of Mendelian inheritance among mice were performed with the GraphPad QuickCalcs data analysis resource (GraphPad Software, Inc., La Jolla, CA, USA) using a χ^2^ test.

#### Cell Lines

Immortalized mouse embryonic palatal mesenchyme (iMEPM) cells were derived from a male *Cdkn2a*^*−/-*^ embryo as previously described (Fantauzzo and Soriano, 2017). iMEPM cells were cultured in medium (Dulbecco’s modified Eagle’s medium [Gibco, Invitrogen] supplemented with 50 U/mL penicillin [Gibco], 50 μg/mL streptomycin [Gibco] and 2 mM L-glutamine [Gibco]) containing 10% FBS (Hyclone Laboratories, Inc.) and grown at 37°C in 5% carbon dioxide.

### METHOD DETAILS

#### RNA-seq and bioinformatics analyses

Secondary palatal shelves (PS) were dissected on ice from three independent biological replicates each of E13.5 *Pdgfra*^*+/+*^ and *Pdgfra*^*PI3K/PI3K*^ embryos and the overlying ectoderm was removed by digestion with 20 mg/mL trypsin for 20 min at 4°C. Maxillary processes (MxP) were dissected on ice from three independent biological replicates each of E11.5 *Srsf3*^*fl/fl*^;*Wnt1-Cre*^*+/+*^ and *Srsf3*^*fl/fl*^;*Wnt1-Cre*^*+/Tg*^ embryos and the overlying ectoderm was removed by digestion with 20 mg/mL trypsin for 15-20 min at 4°C. PS and MxP were subsequently rinsed in 10% fetal bovine serum (FBS) (Hyclone Laboratories, Inc., GE Healthcare, Chicago, IL, USA) for 1 min followed by 1xphosphate buffered saline (PBS). Excess PBS was removed and PS and MxP were submerged in 200 uL RNA*later* stabilization solution (Invitrogen, Carlsbad, CA, USA) for overnight storage at 4°C. The following day excess RNA*later* was removed and the samples were stored at −80°C. Total RNA was simultaneously isolated from all samples using the RNeasy Mini Kit (Qiagen, Inc., Germantown, MD, USA) according to the manufacturer’s instructions. RNA was forwarded to the University of Colorado Anschutz Medical Campus Genomics and Microarray core for quality control, library preparation and sequencing. RNA integrity was assessed on an Agilent 4200 TapeStation using TapeStation Analysis Software A.02.02 (Agilent Technologies, Inc., Santa Clara, CA, USA), revealing RIN^e^ values of 9.6-10.0 for the PS samples and 9.5-10.0 for the MxP samples. RNA concentration was measured with an Infinite M200 PRO Microplate Reader (Tecan Group Ltd., Männedorf, Switzerland). 100 ng of total RNA was used as input into the Universal Plus mRNA-Seq with NuQuant kit (NuGEN, Redwood City, CA, USA). Dual index, stranded libraries were prepared and sequenced on a NovaSeq 6000 Sequencing System using an S4 Flow Cell (Illumina, Inc., San Diego, CA, USA) to an average depth of ~44 million read pairs (2 × 151 bp reads).

BBDuk (from the BBmap v35.85 tool suite) (Bushnell, 2015) was used to perform adapter sequence contamination removal and light quality trimming. For STAR (Dobin et al., 2013) mapping and rMATS (Shen et al., 2014) analysis, read length was standardized to 125 bp to satisfy the input requirements for rMATS. After quality trimming and adapter contamination removal, reads shorter than 125 bp were discarded, and reads longer than 125 bp were cropped to 125 bp. For discovery of AS events, sequence alignment was performed with STAR (v2.7.0) 2-pass mapping, using combined splice junctions reported in the first alignment round. AS events detected via rMATS (v4.0.2, default parameters plus --*readLength 125 --libType fr-secondstrand*) with false discovery rate of ≤ 0.05, a difference in percent spliced in (|Δψ|) ≥ 0.05 and an average count of 10 in either population are reported. For differential expression analysis, RNA sequencing data were quantified at the transcript level via Salmon (v1.1.0) (Patro et al., 2017) using GENCODE (Frankish et al., 2019) vM19 annotation, summarized to gene-level estimated counts via the R package tximport (Soneson et al., 2016) and analyzed for differential expression via DESeq2 (Love et al., 2014). Significant changes in gene-level expression are reported for cases with adjusted p-value of ≤ 0.05. Raw read pairs, trimmed read pairs (125 bp) for STAR input, STAR unique mapping rate, trimmed read pairs for Salmon input and Salmon mapping rate per sample can be found in Table S1. For exons identified to have a skipped exon event, a region of interest (ROI) was generated to include the skipped exon plus 250 bp flanking both ends. ROIs were specific to the strand of the skipped exon. ROIs were analyzed by RBPmap (Paz et al., 2014) to search for Srsf3 (both “cuckucy” and “wcwwc”) motifs using default parameters and the mm10 genome assembly. RBPmap output was parsed and filtered using a custom R script and only exact motif matches were kept. Motifs were summarized by “n” per ROI. Motif location was determined by scoring each motif hit location as upstream, within or downstream of the exon. Statistical analyses of motif density between significant versus non-significant skipped exon events were performed using a Wilcoxon rank sum test with continuity correction and statistical analyses of motif location were performed using a χ^2^ test. To predict potential outcomes of AS events, the R package SPLINTER (Low, 2020) was run with default parameters using the skipped exon ‘JCEC’ rMATS output file. A transcript and coding sequence database was prepared from the GTF file used above for STAR alignment. For each AS event identified by rMATS, SPLINTER identified compatible transcripts and compared the compatible transcripts before and after removal of the exon. One outcome was predicted per each compatible transcript.

#### qPCR

Total RNA was isolated using the RNeasy mini kit (Qiagen, Inc.) according to the manufacturer’s instructions. First strand cDNA was synthesized using a ratio of 2:1 random primers: oligo (dT) primer and SuperScript II RT (Invitrogen) according to the manufacturer’s instructions. All reactions were performed with 1x ThermoPol buffer (0.02 M Tris pH 8.8, 0.01 M KCl, 0.01 M (NH_4_)_2_SO_4_, 2 mM MgSO_4_ and 0.1% Triton X-100), 200 μM dNTPs, 200 nM primers (Integrated DNA Technologies, Inc., Coralville, IA, USA), 0.6 U Taq polymerase and 1 μg cDNA in a 25 μL reaction volume. The primers used can be found in Table S10. The following PCR protocol was used for *Chrd, Cask* and *Smad7*: step 1, 3 min at 94°C; step 2, 30 sec at 94°C; step 3, 30 sec at 50°C; step 4, 30 sec at 72°C; repeat steps 2–4 for 34 cycles; and step 5, 2 min at 72°C. The following PCR protocol was used for *Srsf3* and *Prkd2*: step 1, 3 min at 94°C; step 2, 30 sec at 94°C; step 3, 30 sec at 48°C; step 4, 30 sec at 72°C; repeat steps 2–4 for 34 cycles; and step 5, 2 min at 72°C. The following PCR protocol was used for *Fgfr2*: step 1, 3 min at 94°C; step 2, 30 sec at 94°C; step 3, 30 sec at 49.5°C; step 4, 25 sec at 72°C; repeat steps 2–4 for 34 cycles; and step 5, 2 min at 72°C. The following PCR protocol was used for *B2m*: step 1, 3 min at 95°C; step 2, 30 sec at 95°C; step 3, 30 sec at 56°C; step 4, 30 sec at 72°C; repeat steps 2–4 for 34 cycles; and step 5, 2 min at 72°C. The following PCR protocol was used for *Melk* and *Limk2*: step 1, 3 min at 94°C; step 2, 30 sec at 94°C; step 3, 30 sec at 51°C; step 4, 30 sec at 72°C; repeat steps 2–4 for 34 cycles; and step 5, 2 min at 72°C. The following PCR protocol was used for *Dmpk*: step 1, 3 min at 94°C; step 2, 30 sec at 94°C; step 3, 30 sec at 55°C; step 4, 30 sec at 72°C; repeat steps 2–4 for 34 cycles; and step 5, 2 min at 72°C. Two-thirds of total PCR products were electrophoresed on a 2% agarose/TBE gel containing ethidium bromide and photographed on an Aplegen Omega Fluor Gel Documentation System (Aplegen, Inc., Pleasanton, CA). When applicable, quantifications of band intensity were performed with ImageJ software (version 1.51m9, National Institutes of Health). Statistical analyses were performed with Prism 8 (GraphPad Software, Inc.) using a paired t-test. qPCR reactions were performed using samples from at least three embryos per genotype.

#### In situ *hybridization*

Probe template was amplified by PCR from E13.5 head cDNA using the following primers: 5’-GACTCTAGAGATAAGGGTAGGAACCACAC-3’ and 5’-GAGAAGCTTGGAATGTTTTACCTGGACTTG-3’. PCR product was cloned into the pBluescript II KS(+) dual promoter (T7 and T3) phagemid vector (Agilent Technologies, Inc.) and standard procedures were followed for the preparation of DIG-labeled cRNA (Roche Diagnostics, Indianapolis, IN, USA) anti-sense (AS) and control sense (S) probes. Whole mount *in situ* hybridization was performed based on a previously published protocol (Wilkinson, 1998) using 1 μg/mL probe and anti-DIG-AP Fab fragment primary antibody (1:2,000; Roche Diagnostics). Stained embryos were photographed using an Axiocam 105 color digital camera (Carl Zeiss, Inc.) fitted onto a Stemi 508 stereo microscope (Carl Zeiss, Inc.). *In situ* hybridization experiments were performed simultaneously using anti-sense and sense probes across at least two independent experiments per timepoint.

#### Immunofluorescence analysis

Embryos were fixed in 4% paraformaldehyde (PFA) in 1x PBS and infiltrated with 30% sucrose in PBS before being mounted in O.C.T. compound (Sakura Finetek USA Inc., Torrance, CA, USA). Sections (8 μm) were deposited on glass slides. Sections were fixed in 4% PFA in PBS with 0.1% Triton X-100 for 10 min and washed in PBS with 0.1% Triton-X 100. Sections were blocked for 1 hr in 5% normal donkey serum (Jackson ImmunoResearch Inc., West Grove, PA, USA) in PBS and incubated overnight at 4°C in primary antibody in 1% normal donkey serum in PBS. After washing in PBS, sections were incubated in Alexa Fluor 488-conjugated donkey anti-rabbit secondary antibody (1:1,000; Invitrogen) diluted in 1% normal donkey serum in PBS with 2 μg/mL 4’,6-diamidino-2-phenylindole (DAPI; Sigma-Aldrich Corp., St. Louis, MO, USA) for 1 hr. Sections were mounted in Aqua Poly/Mount mounting medium (Polysciences, Inc., Warrington, PA, USA) and photographed using an Axiocam 506 mono digital camera (Carl Zeiss, Inc., Thornwood, NY, USA) fitted onto an Axio Observer 7 fluorescence microscope (Carl Zeiss, Inc.). The following antibodies were used for immunofluorescence analysis: Srsf3 (1:500; Abcam plc, Cambridge, MA, USA); Ki67 (1:300; Invitrogen). Srsf3 immunofluorescence analysis was performed on multiple sections from individual E8.5-E10.5 embryos and from four E13.5 embryos. All Ki67-positive signals were confirmed by DAPI staining. The percentage of Ki67-positive cells was determined in three embryos per genotype, with up to four sections analyzed per individual embryo. Analyzed sections within a given embryo were 5-10 sections apart, representing a distance of 40-80 μm. Graphed data represent averages from three independent embryos. Statistical analyses were performed on values from individual sections with Prism 8 (GraphPad Software, Inc.) using a two-tailed, unpaired t-test with Welch’s correction.

#### Immunoprecipitations and Western blotting

To induce PDGFRα signaling, passage 23-29 iMEPM cells at ~70% confluence were serum-starved for 24 h in medium containing 0.1% FBS and stimulated with 10 ng/mL PDGF-AA ligand (R&D Systems, Minneapolis, MN, USA) for the indicated length of time. When applicable, cells were pretreated with 50 μM LY294002 (Sigma-Aldrich Corp.) 1 h before PDGF-AA ligand stimulation. Subcellular fractions were generated by harvesting cells in ice-cold PBS, resuspending cells in hypotonic buffer (20 mM Tris-HCl pH 7.4, 10 mM NaCl, 3 mM MgCl_2_ and 0.5% Nonidet P-40) and collecting cytoplasmic lysates by centrifugation at 3,000 rpm at 4°C for 10 min. The nuclear pellet was resuspended in cell extraction buffer (100 mM Tris-HCl pH 7.4, 100 mM NaCl, 10% glycerol, 1% Triton X-100, 1 mM EDTA, 1 mM EGTA, 0.1% SDS, 1x complete Mini protease inhibitor cocktail (Roche Diagnostics), 1 mM PMSF, 1 mM NaF, 2 mM Na_3_VO_4_, 0.5% deoxycholate and 20 mM Na_4_P_2_O_7_) and nuclear lysates collected by centrifugation at 13,000 rpm at 4°C for 30 min. For immunoprecipitations, cell lysates were incubated with magnetic bead-conjugated phospho-Akt substrate primary antibody (1:30; 110B7E; Cell Signaling Technology) overnight at 4°C. Beads were washed with lysis buffer five times and the precipitated proteins were eluted with Laemmli buffer containing 10% β-mercaptoethanol, heated for 5 min at 100°C and separated by SDS-PAGE. Maxillary processes (MxP) were dissected on ice from multiple E11.5 *Srsf3*^*fl/fl*^;*Wnt1-Cre*^*+/+*^ and *Srsf3*^*fl/fl*^;*Wnt1-Cre*^*+/Tg*^ embryos and the overlying ectoderm was removed by digestion with 20 mg/mL trypsin for 15-20 min at 4°C. MxP were subsequently rinsed in 10% FBS (Hyclone Laboratories, Inc.) for 1 min followed by 1x PBS. Excess PBS was removed and MxP were snap frozen in 100% ethanol on dry ice and stored at −80°C. MxP from three embryos per genotype were thawed on ice and pooled to form biological replicates. Protein lysates were generated by resuspending tissues in ice-cold NP-40 lysis buffer (20 mM Tris-HCl pH 8, 150 mM NaCl, 10% glycerol, 1% Nonidet P-40, 2 mM EDTA, 1x complete Mini protease inhibitor cocktail (Roche Diagnostics), 1 mM PMSF, 10 mM NaF, 1 mM Na_3_VO_4_, 25 mM β-glycerophosphate) and collecting cleared lysates by centrifugation at 12,000 rpm at 4°C for 20 min. Laemmli buffer containing 10% β-mercaptoethanol was added to the lysates, which were heated for 5 min at 100°C. Proteins were subsequently separated by SDS-PAGE. Western blot analysis was performed according to standard protocols using horseradish peroxidase-conjugated secondary antibodies. The following antibodies were used for Western blotting: Srsf3 (1:1,000; Abcam plc); Lamin B1 (1:1,000; D4Q4Z; Cell Signaling Technology, Inc.); β-tubulin (1:1,000; E7; Developmental Studies Hybridoma Bank, Iowa City, IA, USA); Phospho-Akt (1:1,000; Ser473; Cell Signaling Technology, Inc.); Akt (1:1,000; Cell Signaling Technology, Inc.); Phospho-VASP (1:1,000; Ser157; Cell Signaling Technology, Inc.); VASP (1:1,000; Cell Signaling Technology, Inc.); horseradish peroxidase-conjugated goat anti-rabbit IgG (1:10,000; Jackson ImmunoResearch Laboratories, Inc., West Grove, PA, USA); horseradish peroxidase-conjugated goat anti-mouse IgG (1:10,000; Jackson ImmunoResearch Laboratories, Inc.). Quantifications of signal intensity were performed with ImageJ software (version 1.51m9, National Institutes of Health). Relative phospho-Srsf3 levels were determined by normalizing to Lamin B1 and β-tubulin for the nuclear and cytoplasmic fractions, respectively. Relative phospho-Vasp levels were determined by normalizing to total Vasp levels. When applicable, statistical analyses were performed with Prism 8 (GraphPad Software, Inc.) using a paired t-test. Immunoprecipitation and Western blotting experiments were performed across at least three independent experiments, with the exception of the phospho-Vasp and Vasp Western blots, which were performed across two independent experiments, each consisting of pooled lysates from three embryos per genotype.

#### Morphological and histological analyses

Embryos were dissected at multiple timepoints (day of plug considered 0.5 days) in 1x PBS and fixed overnight at 4°C in 4% PFA in PBS. Embryos were photographed using an Axiocam 105 color digital camera (Carl Zeiss, Inc., Thornwood, NY, USA) fitted onto a Stemi 508 stereo microscope (Carl Zeiss, Inc.). For histological analyses, following fixation embryos were dehydrated through a graded ethanol series and embedded in paraffin. After deparaffinization and rehydration, sections (8 μm) were stained with hematoxylin and eosin and permanently mounted with Permount (Thermo Fisher Scientific). Sections were photographed using an Axiocam 503 color digital camera (Carl Zeiss, Inc.) fitted onto an Axio Observer 7 fluorescence microscope (Carl Zeiss, Inc.). One representative embryo per genotype per timepoint was photographed for morphological analysis and one representative embryo per genotype per timepoint was subjected to histological analysis.

#### Whole-mount DAPI staining

Whole-mount DAPI staining was performed according to a previously published protocol (Sandell et al., 2012), with the exception that staining was performed with 10 μg/mL DAPI (Sigma-Aldrich Corp.) for 1 hr at room temperature. Embryos were photographed using an Axiocam 506 mono digital camera (Carl Zeiss, Inc.) fitted onto an Axio Observer 7 fluorescence microscope (Carl Zeiss, Inc.). Extended Depth of Focus was applied to z-stacks using ZEN Blue software (Carl Zeiss, Inc.) to generate images with the maximum depth of field. One representative embryo per genotype per timepoint not possessing the *ROSA26*^*mTmG*^ allele was photographed for E10.5-E14.5 embryos. At least three embryos per genotype per timepoint possessing the *ROSA26*^*mTmG*^ allele were photographed for E8.0-E10.5 embryos.

#### Skeletal preparations

E18.5 embryos were skinned, eviscerated, fixed in 95% ethanol and stained in 0.015% Alcian blue, 0.005% Alizarin red and 5% glacial acetic acid in 70% ethanol at 37°C. Embryos were then cleared in 1% KOH and transferred to solutions of decreasing KOH concentration and increasing glycerol concentration. Skeletons were photographed using an Axiocam 105 color digital camera (Carl Zeiss, Inc.) fitted onto a Stemi 508 stereo microscope (Carl Zeiss, Inc.). At least three embryos per genotype were assayed.

#### TUNEL assay

Sections (8 μm) of PFA-fixed, sucrose-infiltrated, O.C.T.-mounted embryos were deposited on glass slides. Apoptotic cells were identified using the *In Situ* Cell Death Detection Kit, Fluorescein (Sigma-Aldrich Corp.) according to the manufacturer’s instructions for the treatment of cryopreserved tissue sections. Sections were mounted in VECTASHIELD® Antifade Mounting Medium with DAPI (Vector Laboratories, Burlingame, CA, USA) and photographed using an Axiocam 506 mono digital camera (Carl Zeiss, Inc.) fitted onto an Axio Observed 7 fluorescence microscope (Carl Zeiss, Inc.). All positive signals were confirmed by DAPI staining. The percentage of TUNEL-positive cells was determined in three embryos per genotype, with up for four sections analyzed per individual embryo. Analyzed sections wtihin a given embryo were 5-10 sections apart, representing a distance of 40-80 μm. Graphed data represent averages from three independent embryos. Statistical analyses were performed on values from individual sections with Prism 8 (GraphPad Software, Inc.) using a two-tailed, unpaired t-test with Welch’s correction.

#### qRT-PCR

Total RNA was isolated and cDNA was synthesized as above. qRT-PCR was performed on a CFX Connect Real-Time PCR Detection System and analyzed with CFX Manager software (version 3.1; Bio-Rad Laboratories, Inc., Hercules, CA, USA). All reactions were performed with SYBR Select Master Mix (Applied Biosystems, Foster City, CA, USA), 300 nM primers (Integrated DNA Technologies, Inc.) and cDNA in a 20 μL reaction volume. The primers used can be found in Table S10. The following PCR protocol was used: step 1, 2 min at 50°C; step 2, 2 min at 95°C; step 3, 15 sec at 95°C; step 4, 1 min at 60°C; repeat steps 3 and 4 for 39 cycles; step 5 (melting curve), 5 sec per 0.5°C increment from 65°C to 95°C. All samples were run in triplicate and normalized against an endogenous internal control, *B2m*. Statistical analyses were performed with Prism 8 (GraphPad Software, Inc.) using a paired t-test. qRT-PCR reactions were performed using samples from three embryos per genotype.

### QUANTIFICATION AND STATISTICAL ANALYSIS

AS events detected by rMATS were considered significant with a false discovery rate of ≤ 0.05, a difference in percent spliced in (|Δψ|) ≥ 0.05 and an average count of 10 in either population analyzed. Differential gene expression detected by DESeq2 was considered significant with an adjusted p-value of ≤ 0.05. Statistical analyses of motif density detected by RBPmap between significant versus non-significant skipped exon events were performed using a Wilcoxon rank sum test with continuity correction and statistical analyses of motif location were performed using a χ^2^ test. Gene ontology analysis was performed with various libraries from the Enrichr gene list enrichment analysis tool (Chen et al., 2013; Kuleshov et al., 2016) and terms with p < 0.05 were considered significant. For qPCR analyses, quantifications of band intensity were performed with ImageJ software (version 1.51m9, National Institutes of Health) and statistical analyses were performed with Prism 8 (GraphPad Software, Inc.) using a paired t-test. Statistical analyses of Mendelian inheritance among mice were performed with the GraphPad QuickCalcs data analysis resource (GraphPad Software, Inc.) using a χ^2^ test. For Ki67 immunofluorescence and TUNEL analyses, statistical analyses were performed on values from individual sections across three independent embryos per genotype with Prism 8 (GraphPad Software, Inc.) using a two-tailed, unpaired t-test with Welch’s correction. For qRT-PCR analyses, Cq values were determined using CFX Manager software (version 3.1; Bio-Rad Laboratories, Inc.) and statistical analyses were performed with Prism 8 (GraphPad Software, Inc.) using a paired t-test. For Western blot analyses, quantifications of band intensity were performed with ImageJ software (version 1.51m9, National Institutes of Health) and statistical analyses were performed with Prism 8 (GraphPad Software, Inc.) using a paired t-test. Bar graph data are presented with mean values. Scatter dot plot and line graph data are presented as mean ± SEM. Unless otherwise noted, comparisons with a p-value < 0.05 were considered significant in all statistical analyses.

### DATA AND CODE AVAILABILITY

The RNA-sequencing datasets generated during this study are available at the Gene Expression Omnibus (GEO) accession code GSE161510.

**Supplemental Item Titles (for items not included in Supplemental Information file) Table S2, related to Figure 1.** rMATS output for *Pdgfra*^*+/+*^ versus *Pdgfra*^*PI3K/PI3K*^ RNA-seq analysis. Skipped exon (SE) events additionally contain RBPmap output for Srsf3.

**Table S4, related to Figure 1.** DESeq2 output for *Pdgfra*^*+/+*^ versus *Pdgfra*^*PI3K/PI3K*^ RNA-seq analysis.

**Table S6, related to Figure 6.** DESeq2 output for *Srsf3*^*fl/fl*^;*Wnt1-Cre*^*+/+*^ versus *Srsf3*^*fl/fl*^;*Wnt1-Cre*^*+/Tg*^ RNA-seq analysis.

**Table S8, related to Figure 7.** rMATS output for *Srsf3*^*fl/fl*^;*Wnt1-Cre*^*+/+*^ versus *Srsf3*^*fl/fl*^;*Wnt1-Cre*^*+/Tg*^ RNA-seq analysis. Skipped exon (SE) events additionally contain RBPmap output for Srsf3 and SPLINTER output.

