## Supplemental Information for "Srsf3 mediates alternative RNA splicing downstream of PDGFRα signaling"

### Supplemental Tables

**Table S1, related to Figures 1, 6 and 7.** RNA-seq sample information.

| Sample | Raw read pairs | Trimmed read pairs (125 bp) for STAR input | STAR unique mapping rate | Trimmed read pairs for Salmon input | Salmon mapping rate |
|---|---|---|---|---|---|
| WT_1 | 34222771 | 27870420 | 86.71 | 33922569 | 90.37 |
| WT_2 | 39737843 | 32252735 | 86.49 | 39393855 | 89.68 |
| WT_3 | 38588717 | 34454350 | 87.45 | 38228994 | 88.28 |
| MUT_1 | 34129241 | 29490238 | 87.67 | 33845444 | 90.14 |
| MUT_2 | 36883110 | 31631761 | 87.66 | 36506989 | 88.51 |
| MUT_3 | 42198499 | 36007926 | 87.27 | 41818779 | 89.45 |
| Control_1 | 41471062 | 28063811 | 88.15 | 41450695 | 90.67 |
| Control_2 | 42396709 | 29548001 | 87.39 | 42352348 | 90.87 |
| Control_3 | 53748421 | 37788049 | 87.23 | 53718575 | 89.79 |
| cKO_1 | 48542139 | 34411636 | 87.29 | 48519668 | 89.05 |
| cKO_2 | 70099132 | 48039070 | 86.37 | 70051229 | 90.39 |
| cKO_3 | 45248549 | 31400746 | 87.25 | 45211149 | 90.30 |

WT = *Pdgfra*$^{+/+}$; MUT = *Pdgfra*$^{PI3K/PI3K}$; Control = *Srsf3*$^{fl/fl}$;*Wnt1-Cre*$^{+/+}$; cKO = *Srsf3*$^{fl/fl}$;*Wnt1-Cre*$^{+/Tg}$

**Table S3, related to Figure 1.** Gene name, exon coordinates and ∆PSI values from *Pdgfra*$^{+/+}$ versus *Pdgfra*$^{PI3K/PI3K}$ RNA-seq analysis for skipped exon events with one or more Srsf3 motifs and a corresponding mouse model with a craniofacial phenotype. Mouse models with a cleft secondary palate are highlighted in bold.

| Gene | Exon start | Exon end | ∆PSI |
|---|---|---|---|
| *Gng8* | chr7:16893120 | chr7:16893219 | 36.1 |
| *Vdr* | chr15:97869317 | chr15:97869423 | 25.4 |
| *Gng8* | chr7:16892449 | chr7:16892511 | 20.8 |
| *Akap10* | chr11:61904774 | chr11:61904859 | 19.9 |
| *Ptk2* | chr15:73365040 | chr15:73365195 | 18.4 |
| *Nf1* | chr11:79463232 | chr11:79463295 | 18.0 |
| *Bcas3* | chr11:85822021 | chr11:85822112 | 17.5 |
| ***Fuz*** | chr7:44896546 | chr7:44896645 | 16.7 |
| ***Fuz*** | chr7:44896546 | chr7:44896668 | 15.4 |
| ***Chrd*** | chr16:20739953 | chr16:20740056 | 14.3 |
| ***Fuz*** | chr7:44896914 | chr7:44896999 | 13.1 |
| ***Cask*** | chrX:13552372 | chrX:13552441 | 11.3 |
| *Vps54* | chr11:21268784 | chr11:21268863 | 10.2 |
| ***Smad7*** | chr18:75375873 | chr18:75375948 | 9.5 |
| ***Qrich1*** | chr9:108518766 | chr9:108518826 | 8.2 |
| *Sptan1* | chr2:30001099 | chr2:30001159 | 7.8 |
| *Dyrk1a* | chr16:94611418 | chr16:94611504 | 7.7 |
| *E2f5* | chr3:14602647 | chr3:14602712 | 6.5 |
| *Zeb2* | chr2:45017396 | chr2:45017465 | 6.3 |
| ***Map3k7*** | chr4:31994873 | chr4:31994954 | 6.2 |
| *Apaf1* | chr10:91023719 | chr10:91023848 | 5.6 |
| ***Tgfb2*** | chr1:186690720 | chr1:186690804 | 5.2 |
| *Inppl1* | chr7:101823512 | chr7:101823646 | -5.7 |
| *Pdpk1* | chr17:24107228 | chr17:24107271 | -12.6 |
| ***Dzip1l*** | chr9:99632636 | chr9:99632748 | -14.4 |

**Table S5, related to Figure 4.** Progeny from *Srsf3*$^{fl/fl}$ x *Srsf3*$^{fl/fl}$*;Wnt1-Cre*$^{+/Tg}$ crosses.

| Age | *Srsf3*$^{+/fl}$*; W1C*$^{+/+}$ | *Srsf3*$^{+/fl}$*; W1C*$^{+/Tg}$ | *Srsf3*$^{fl/fl}$*; W1C*$^{+/+}$ | *Srsf3*$^{fl/fl}$*;W1C*$^{+/Tg}$ | | | | Chi-square* |
|---|---|---|---|---|---|---|---|---|
| | | | | Total | Live | Dead but developmentally equal | Resorbed/ delayed | |
| **E8.5** | 6 | 4 | 11 | 7 | 7 (25.0%) | 0 | 0 | p = 1.0000 |
| **E9.5** | 13 | 9 | 11 | 9 | 9 (21.4%) | 0 | 0 | p = 0.5930 |
| **E10.5** | 35 | 32 | 32 | 34 | 30 (22.6%) | 1 | 3 | p = 0.7613 |
| **E11.5** | 32 | 26 | 18 | 22 | 20 (20.4%) | 1 | 1 | p = 0.4460 |
| **E12.5** | 12 | 15 | 20 | 10 | 6 (10.5%) | 1 | 3 | p = 0.0411 |
| **E14.5** | 5 | 5 | 2 | 2 | 1 (7.1%) | 0 | 1 | p = 0.1495 |
| **E15.5** | 14 | 8 | 13 | 7 | 0 (0%) | 0 | 7 | p = 0.0006 |
| **E18.5** | 9 | 8 | 7 | 3 | 1 (3.7%) | 2 | 0 | p = 0.0956 |
| **P0** | 18 | 14 | 20 | 2 | 0 (0%) | 2 | 0 | p = 0.0003 |

*Chi square analysis was performed with live and dead but developmentally equal counts for *Srsf3*$^{fl/fl}$*;Wnt1-Cre*$^{+/Tg}$ embryos

**Table S7, related to Figure 6.** Gene name and log2(fold change) values from *Srsf3*$^{fl/fl}$*;Wnt1-Cre*$^{+/+}$ versus *Srsf3*$^{fl/fl}$*;Wnt1-Cre*$^{+/Tg}$ RNA-seq analysis for differentially-expressed genes with a corresponding mouse model with a craniofacial phenotype. Mouse models with a cleft secondary palate are highlighted in bold.

| Gene | log2(fold change) |
|---|---|
| ***Sim2*** | 4.15812767 |
| *Ascl1* | 3.90697996 |
| *Emx2* | 3.27243847 |
| *Grem2* | 2.99306483 |
| *Thrb* | 2.85205581 |
| *Smoc2* | 2.74134256 |
| ***Inhba*** | 2.52841522 |
| *Aldh1a2* | 2.42461884 |
| ***Kcnj2*** | 2.28766851 |
| *Capn6* | 1.39714305 |
| ***Eya4*** | 1.39646025 |
| *Alx1* | 1.17541114 |
| *Slc8a1* | 1.17422346 |
| *L1cam* | 1.16812077 |
| *Eda* | 1.12211529 |
| *Lgals1* | 0.99776335 |
| *Gata3* | 0.99406115 |
| ***Foxd3*** | 0.98552765 |
| ***Pcsk5*** | 0.81156592 |
| ***Fgf10*** | 0.80927165 |
| *Alx4* | 0.72872988 |
| *Bmp1* | 0.59722314 |
| *Irx3* | 0.58022535 |
| ***Gli2*** | 0.55774872 |
| *Cdon* | 0.54318469 |
| *Sulf2* | 0.51966198 |
| *Lima1* | 0.48945473 |
| ***Pdgfc*** | -0.6367654 |
| ***Piga*** | -0.6485028 |
| *Col18a1* | -0.6782651 |
| *Notch1* | -0.7699911 |
| *Cdh5* | -0.7887263 |
| *Etv5* | -0.7957678 |
| ***Col2a1*** | -0.8217745 |
| *Lyn* | -0.9220268 |
| *Fbn2* | -0.9424215 |
| *Klf2* | -0.9430508 |
| ***Hspg2*** | -0.9611855 |
| ***Jag2*** | -0.9686243 |
| *Kdr* | -0.985219 |

| | |
|---|---|
| ***Fras1*** | -1.0847412 |
| *Cxxc5* | -1.2326119 |
| *Tfpi* | -1.2517335 |
| *Eng* | -1.2579656 |
| ***Tbx1*** | -1.5287566 |
| *Lama5* | -1.5415792 |
| ***Ctgf*** | -1.6336439 |
| *Tcf15* | -1.8120891 |
| ***Trp63*** | -1.9570749 |
| *Frem2* | -2.3166877 |
| *Kirrel3* | -2.3488613 |
| ***Hand2*** | -3.0639499 |
| *Pou3f4* | -3.2610681 |
| *Lin28a* | -3.3688949 |
| *Alx3* | -3.4020552 |
| *Foxc1* | -3.434487 |
| *St14* | -4.1221467 |
| *Cyp26c1* | -4.8790515 |
| *Zic5* | -5.6682824 |
| ***Pitx1*** | -5.9498278 |
| ***Kynu*** | -6.1435873 |
| *Foxg1* | -7.343262 |
| *Zic2* | -7.3466391 |
| *Hmx1* | -7.9709667 |
| ***Zic3*** | -9.4878579 |

**Table S9, related to Figure 7.** Gene name, exon coordinates and ∆PSI values from *Srsf3*$^{fl/fl}$*;Wnt1-Cre*$^{+/+}$ versus *Srsf3*$^{fl/fl}$*;Wnt1-Cre*$^{+/Tg}$ RNA-seq analysis for skipped exon events with one or more Srsf3 motifs and a corresponding mouse model with a craniofacial phenotype. Mouse models with a cleft or arched secondary palate are highlighted in bold.

| Gene | Exon start | Exon end | ∆PSI |
|---|---|---|---|
| *Ptk2* | chr15:73365040 | chr15:73365195 | 41.8 |
| *Kars* | chr8:112010221 | chr8:112010280 | 38.5 |
| *Bcl3* | chr7:19809141 | chr7:19809259 | 16.5 |
| ***Tbc1d32*** | chr10:56161134 | chr10:56161217 | 13.7 |
| ***Golgb1*** | chr16:36886130 | chr16:36886228 | 13.1 |
| ***Slc35d1*** | chr4:103208109 | chr4:103208212 | 12.0 |
| *Slc4a4* | chr5:89232763 | chr5 89232860 | 11.4 |
| *B3glct* | chr5:149709319 | chr5:149709429 | 10.4 |
| ***Ncor2*** | chr5:125019818 | chr5:125020046 | 10.3 |
| *Evc* | chr5:37314761 | chr5:37314827 | 10.1 |
| ***Ncoa6*** | chr2:155405474 | chr2:155408450 | 9.5 |
| *Ltbp1* | chr17:75315002 | chr17:75315125 | 9.0 |
| *Ehmt1* | chr2:24863771 | chr2:24863908 | 8.8 |
| *Gnas* | chr2:174328107 | chr2:174330434 | 8.6 |
| ***Ncor2*** | chr5:125124769 | chr5:125124980 | 8.3 |
| *Tcirg1* | chr19:3897467 | chr19:3897681 | 8.2 |
| *Rps6ka3* | chrX:159278109 | chrX:159278203 | 7.9 |
| *Ltbp1* | chr17:75352686 | chr17:75352830 | 7.8 |
| *Agap1* | chr1:89789162 | chr1:89789324 | 7.6 |
| ***Fuz*** | chr7:44896914 | chr7:44896999 | 7.4 |
| *Atpaf2* | chr11:60411666 | chr11:60411729 | 7.0 |
| *Large1* | chr8:72912055 | chr8:72912160 | 6.7 |
| *Casp9* | chr4:141796437 | chr4:141796834 | 6.0 |
| *Casp9* | chr4:141796434 | chr4:141796834 | 5.5 |
| *Napa* | chr7:16112583 | chr7:16112630 | 5.3 |
| *Nr2c2* | chr6:92152314 | chr6:92152417 | 5.2 |
| ***Trps1*** | chr15:50863882 | chr15:50863927 | -6.5 |
| *Otulin* | chr15:27623281 | chr15:27623358 | -6.8 |
| *B3glct* | chr5:149725410 | chr5:149725487 | -7.3 |
| ***Nprl3*** | chr11:32255438 | chr11:32255568 | -7.7 |
| *Mink1* | chr11:70609553 | chr11:70609717 | -7.7 |
| *Anapc15* | chr7:101897732 | chr7:101897817 | -8.6 |
| *Rhobtb3* | chr13:75917692 | chr13:75917847 | -8.7 |
| ***Tgds*** | chr14:118127494 | chr14:118127585 | -9.1 |
| ***Dnmt3b*** | chr2:153650784 | chr2:153650899 | -9.3 |
| *Mtf2* | chr5:108080824 | chr5:108080841 | -10.2 |
| *Mecom* | chr3:29956305 | chr3:29956489 | -10.2 |
| *Uros* | chr7:133691085 | chr7:133691166 | -10.3 |
| ***Nsd2*** | chr5:33821658 | chr5:33821728 | -10.5 |

| | | | |
|---|---|---|---|
| *App* | chr16:85040250 | chr16:85040307 | -11.1 |
| *Limk2* | chr11:3392916 | chr11:3393016 | -11.5 |
| *Clcn5* | chrX:7318424 | chrX:7318488 | -11.5 |
| *Ltbp3* | chr19:5753928 | chr19:5754048 | -12.5 |
| *Arhgap29* | chr3:121973878 | chr3:121974115 | -12.6 |
| ***Tbx3*** | chr5:119675242 | chr5:119675302 | -14.7 |
| *Clcn5* | chrX:7318383 | chrX:7318488 | -15.7 |
| ***Dzip1l*** | chr9:99632636 | chr9:99632748 | -16.4 |
| *Rps6ka3* | chrX:159278109 | chrX:159278203 | -22.5 |
| ***Zfp950*** | chr19:61118226 | chr19:61118347 | -23.8 |
| *Disp1* | chr1:183202951 | chr1:183203039 | -29.5 |
| *Mbtd1* | chr11:93891338 | chr11:93891388 | -38.0 |

**Table S10, related to Figures 1, 6 and 7.** Primers used in qPCR and qRT-PCR analyses.

| Transcript | Forward primer (5' to 3') | Reverse primer (5' to 3') |
|---|---|---|
| qPCR primers | | |
| *Chrd* | GCTATTTTGATGGTGACCGG | CAGCTCTGATTCTCTGGGAA |
| *Cask* | CAGTTTACCCTGCCACCAGT | CCTTCAAGATTGTGCCAAGC |
| *Smad7* | CTCCTCCTTACTCCAGATAC | CATTCCCCTGAGGTAGATCA |
| *Srsf3* | GAGACCGAGAATCTGTAGGA | CGGGAAAAGCTTCTCCTTCT |
| *Fgfr2* | CTCGGGGATAAATAGCTCCA | CTTCTTGGTCGTGGTCTTCA |
| *Fgfr2* | CTGCATGGTTGACAGTTCTG | CTTCTTGGTCGTGGTCTTCA |
| *B2m* | ACTGACCGGCCTGTATGCTA | TGAAGGACATATCTGACATCTCTA |
| *Melk* | GATGTTTGGAGCATGGGCAT | GCGGTGATGTACAGAAAGCT |
| *Limk2* | GTGGTCCTTCCTGTGTTGTC | CCCATGGCAGAATTCTCCAA |
| *Dmpk* | GTATTAGTGAAAGGGGACCG | CATTCCATCAGGCTGCAGTT |
| *Prkd2* | GACTCCTGAGAAGGTATTCG | CAGGTTGTTGATGAGGTCGA |
| qRT-PCR primers | | |
| *Pdgfc* | CTGCTCAACAATGCTGTGAC | CCTCTTCCTTGAGGAGATTC |
| *Col2a1* | CGCTACACTCAAGTCACTGA | CGGTCTCCATGTTGCAGAAA |
| *Kdr* | GTTTGCCTGGCGATTTTCTC | CAGAAGATACTGTCACCACC |
| *Foxd3* | CCTCTACCCAATCCTGGACT | CACAGTACTGAGAACACAGG |
| *Zic5* | CAAGGCCAAGTACAAGCTCA | GTGGACATGGGAGTGTTTCT |
| *Pou3f4* | CAGGACCACTCTGATGAAGA | GAAGCTCAGTTGTAAGGCCT |
| *B2m* | ACTGACCGGCCTGTATGCTA | TGAAGGACATATCTGACATCTCTA |

### Supplemental Figure Legends

**Figure S1, related to Figure 3.** Total Srsf3 protein levels in the nucleus do not change in response to PDGF-AA ligand treatment. (A) Western blot analysis of total Srsf3 levels in nuclear fractions of iMEPM cells that were untreated or treated with PDGF-AA ligand in a time course analysis from 2 to 240 minutes. WB, Western blot. (B) Line graph depicting quantification of band intensities from three independent experiments as in (A). Data are presented as mean ± SEM. *, $p < 0.05$.

**Figure S2, related to Figure 4.** Efficient deletion of *Srsf3* exons 2 and 3 and maintenance of alternative RNA splicing in *Srsf3* conditional knock-out embryos. (A) Conditional, floxed *Srsf3* locus before (top) and after (bottom) Cre exposure. Boxes represent exons. Colors correspond to protein domains in (B). Black dots indicate sites of forward (F) and reverse (R) primers used in (C). Triangles represent loxP sites. Dotted line surrounds exon 4, which contains a premature termination codon and is excluded from the major transcript. (B) Srsf3 wild-type (top) and conditional knock-out (bottom) proteins. Amino acid residues at the boundaries of the RNA recognition motif (RRM) and arginine/serine-rich (RS) domain are indicated. The epitope for the antibody used in Figure 2, Figure 3 and Figure S1 is indicated. (C) Representative qPCR product gel depicting *Srsf3* transcripts in the facial processes (left) and limb buds (right) of E12.5 $Srsf3^{fl/fl};Wnt1\text{-}Cre^{+/+}$ versus $Srsf3^{fl/fl};Wnt1\text{-}Cre^{+/Tg}$ embryos. (D) Representative qPCR product gel depicting *Fgfr2* and control *B2m* transcripts in the facial processes of E12.5 $Srsf3^{fl/fl};Wnt1\text{-}Cre^{+/+}$ versus $Srsf3^{fl/fl};Wnt1\text{-}Cre^{+/Tg}$ embryos.

**Figure S3, related to Figure 4**. Craniofacial morphological defects in *Srsf3* conditional knock-out embryos. (A-D''') Hematoxylin and eosin-stained coronal sections of E10.5 (A,B) and E12.5 (C-D''') $Srsf3^{fl/fl};Wnt1\text{-}Cre^{+/+}$ embryos (top) and $Srsf3^{fl/fl};Wnt1\text{-}Cre^{+/Tg}$ embryos (bottom). Sections in C-D''' move from anterior to posterior craniofacial structures. LNP, lateral nasal process; MNP, medial nasal process; MxP, maxillary process; MdP, mandibular process, LV, lateral ventricle, NS, nasal septum, TV, third ventricle, PS, secondary palatal shelf, T, tongue. Bars, 1 mm.

**Figure S4, related to Figure 6.** *Srsf3* conditional knock-out RNA-seq reads skip exons 2 and 3. IGV snapshots of *Srsf3* transcripts from RNA-seq analysis of three biological replicates each of E11.5 $Srsf3^{fl/fl};Wnt1\text{-}Cre^{+/+}$ (control) versus $Srsf3^{fl/fl};Wnt1\text{-}Cre^{+/Tg}$ (cKO) maxillary process mesenchyme**.**

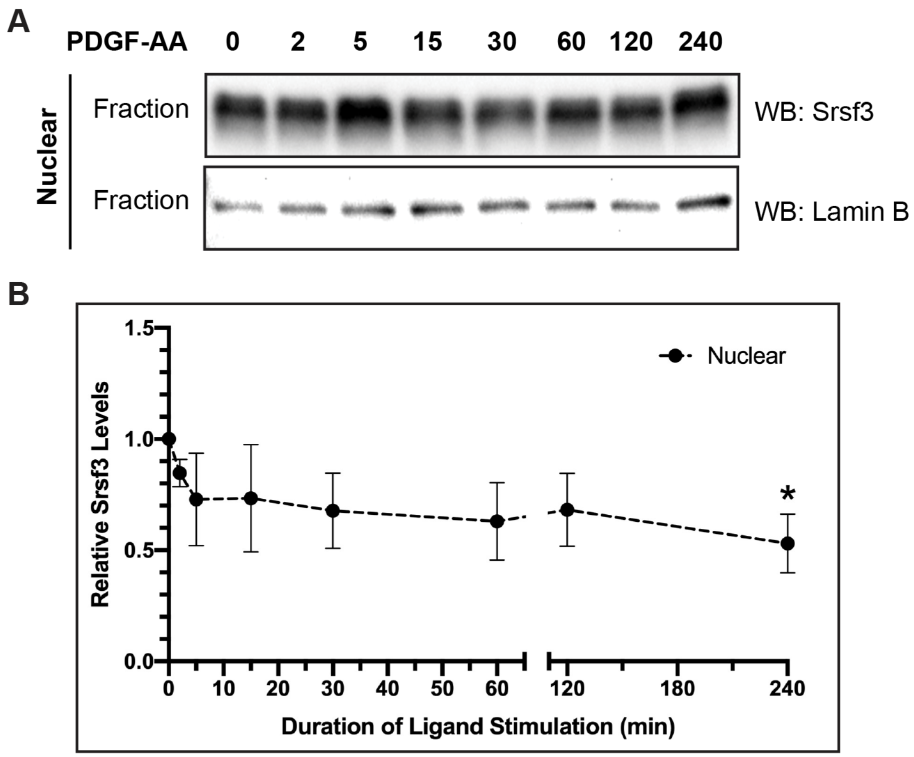
A
PDGF-AA 0 2 5 15 30 60 120 240
Nuclear
Fraction
Fraction
WB: Srsf3
WB: Lamin B
B
Nuclear
Relative Srsf3 Levels
1.5
1.0
0.5
0.0
0 10 20 30 40 50 60 120 180 240
Duration of Ligand Stimulation (min)
*

**Figure S1.**

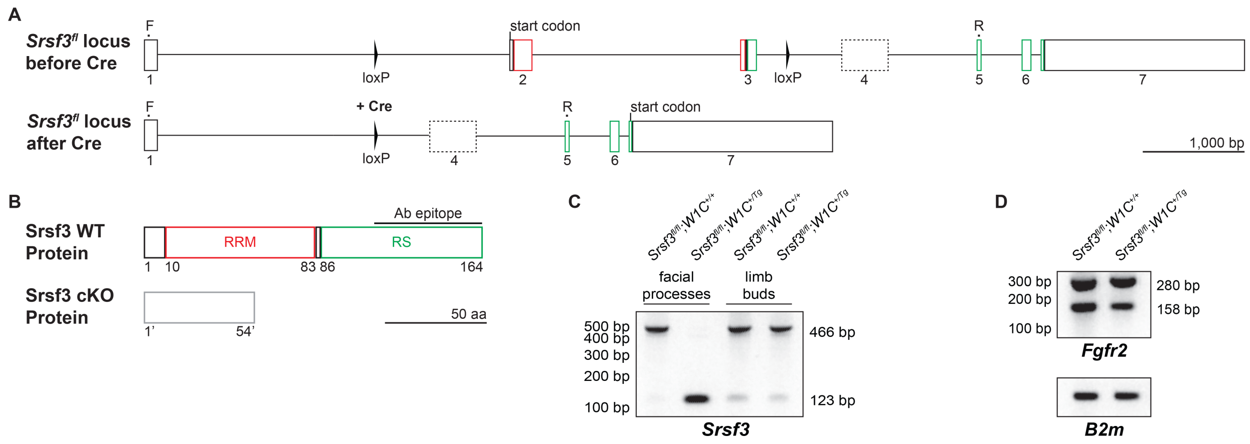
A
Srsf3fl locus before Cre
F
start codon
loxP
R
1
2
3
4
5
6
7
Srsf3fl locus after Cre
+ Cre
start codon
1,000 bp
B
Srsf3 WT Protein
Ab epitope
RRM
RS
1
10
83
86
164
Srsf3 cKO Protein
1'
54'
50 aa
C
facial processes
limb buds
500 bp
400 bp
300 bp
200 bp
100 bp
466 bp
123 bp
Srsf3
D
300 bp
200 bp
100 bp
280 bp
158 bp
Fgfr2
B2m

**Figure S2.**

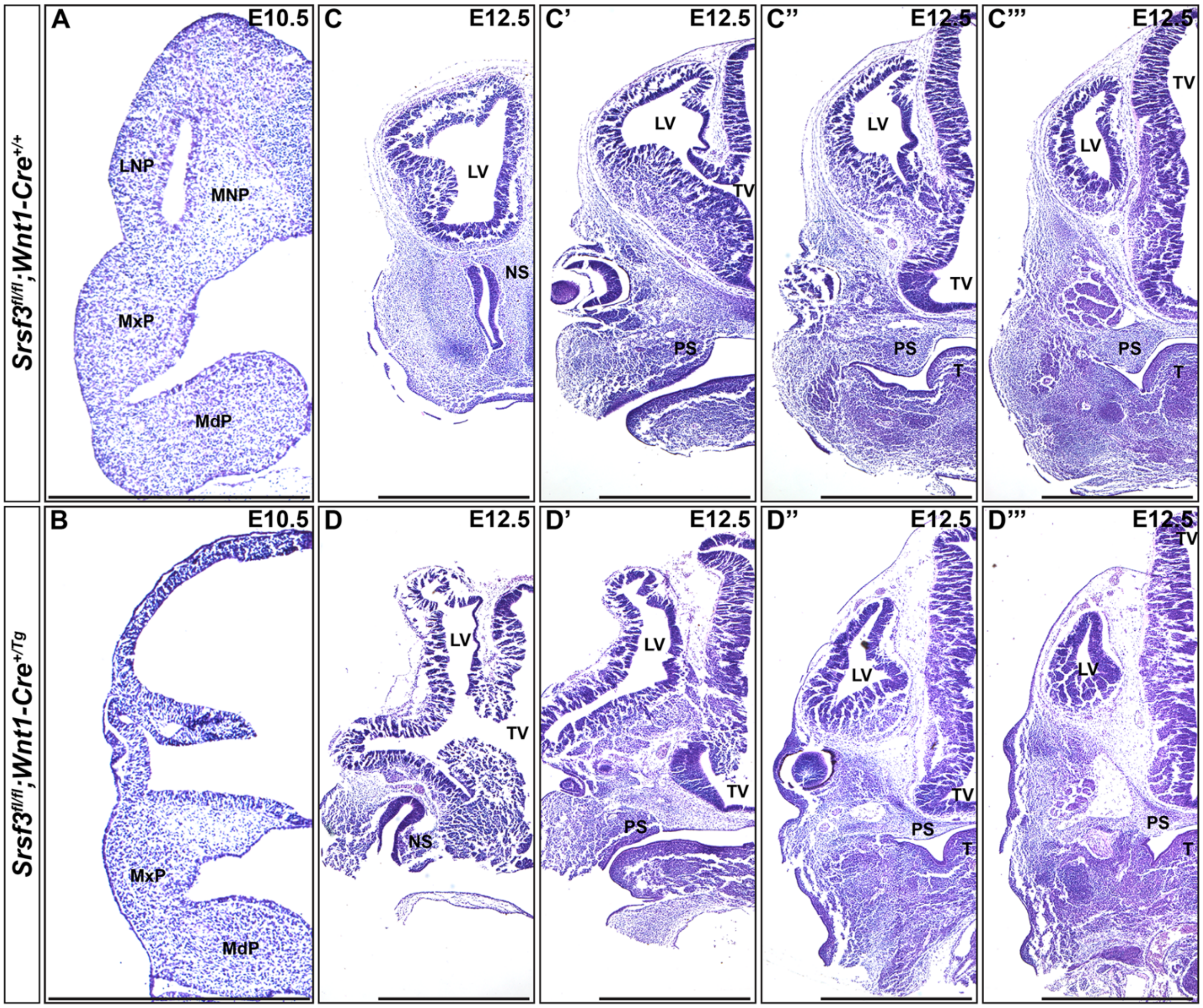

Srsf3fl/fl;Wnt1-Cre+/+
Srsf3fl/fl;Wnt1-Cre+/Tg
A
E10.5
LNP
MNP
MxP
MdP
C
E12.5
LV
NS
C'
E12.5
LV
TV
PS
C''
E12.5
LV
TV
PS
T
C'''
E12.5
TV
LV
PS
T
B
E10.5
MxP
MdP
D
E12.5
LV
TV
NS
D'
E12.5
LV
TV
PS
D''
E12.5
LV
TV
PS
T
D'''
E12.5
TV
LV
PS
T


**Figure S3.**

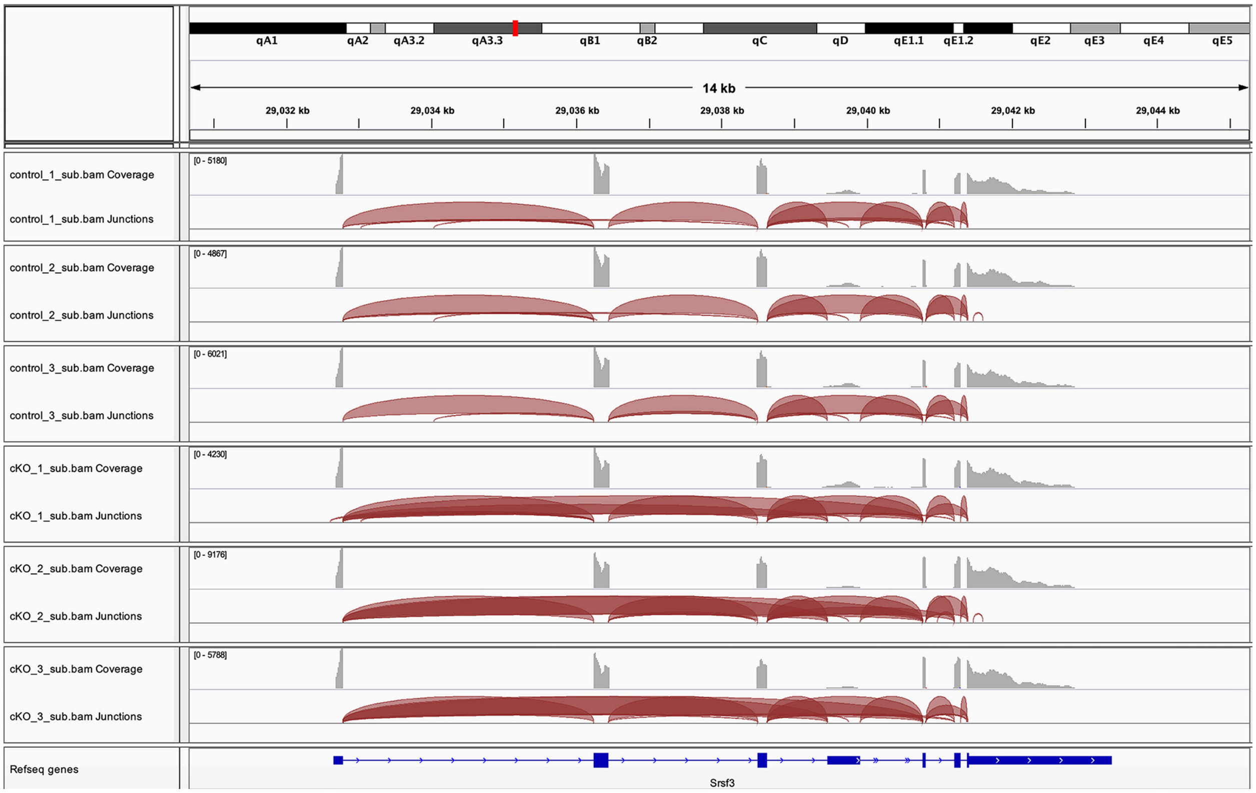
qA1
qA2
qA3.2
qA3.3
qB1
qB2
qC
qD
qE1.1
qE1.2
qE2
qE3
qE4
qE5
14 kb
29,032 kb
29,034 kb
29,036 kb
29,038 kb
29,040 kb
29,042 kb
29,044 kb
control_1_sub.bam Coverage
[0 - 5180]
control_1_sub.bam Junctions
control_2_sub.bam Coverage
[0 - 4867]
control_2_sub.bam Junctions
control_3_sub.bam Coverage
[0 - 6021]
control_3_sub.bam Junctions
cKO_1_sub.bam Coverage
[0 - 4230]
cKO_1_sub.bam Junctions
cKO_2_sub.bam Coverage
[0 - 9176]
cKO_2_sub.bam Junctions
cKO_3_sub.bam Coverage
[0 - 5788]
cKO_3_sub.bam Junctions
Refseq genes
Srsf3

**Figure S4.**
